# Massage Engages Spinal Oxytocin Circuits to Reduce Pain and Enhance Well-Being

**DOI:** 10.64898/2026.01.11.698886

**Authors:** Manon Bohic, Paula Celeste Salamone, Wanhong Zuo, Ahmed Negm, Sasha Leilani Fulton, Shibin Du, Swathi Jayakumar, Jessica Keating, Vanessa Soubeyre, Mark Andrew Gradwell, Aman Upadhyay, Lucero Shorter, Joel Kim, Yukiko Ueno Inoue, Takayoshi Inoue, Brett Mensh, Max Alexander Tischfield, Jiang-Hong Ye, Cedric Peirs, Gaetan Poulen, Nicolas Lonjon, Florence Vachiery-Lahaye, Luc Bauchet, Emmanuel Bourinet, Yuan-Xiang Tao, Hakan Olausson, Ishmail Abdus-Saboor, Rebecca Boehme, Victoria E. Abraira

## Abstract

Context determines how the nervous system interprets sensory information to guide adaptive behavior. Nowhere is this more consequential than in the somatosensory system where the same touch can signal threat from a predator or safety from a caregiver, injury during attack or comfort during healing. Oxytocin, a neuropeptide released across a broad range of social interactions, is well positioned to convey social context to somatosensory circuits, but at what stage of processing does this convergence occur and through what mechanism? Here we show that affiliative touch triggers oxytocin release at the earliest stage of sensory processing, the spinal cord, shaping both pain relief and touch reward through a single, state-dependent circuit that is conserved from mice to humans. In humans, massage drives endogenous oxytocin release, enhanced pain relief and well-being. Modeling this in mice shows that social touch activates hypothalamic oxytocin neurons that project to the spinal cord, where oxytocin signaling is both necessary and sufficient for pain suppression and touch reward. Moreover, spinal calcium imaging and slice electrophysiology demonstrate that oxytocin acts on both excitatory and inhibitory spinal neurons to differentially modulate the relative activity of spinal ascending systems that convey both pain and social touch to the brain. We find that this circuit architecture is conserved in humans, as oxytocin receptors are similarly expressed on spinal excitatory and inhibitory neurons. Massage-induced oxytocin release in humans correlates with slowed somatosensory processing within the spinal cord in response to aversive stimuli and reduced negative affect. Thus, spinal oxytocin circuits provide an evolutionarily conserved mechanism through which affiliative touch reduces pain and enhances well-being, linking the earliest stage of sensory processing to the social bonds on which survival depends.

**Highlights:**

- Massage drives endogenous oxytocin release that correlates with pain relief and enhanced well-being in humans
- Social touch activates hypothalamic oxytocin neurons that project directly to the spinal cord dorsal horn
- Spinal oxytocin signaling is both necessary and sufficient for pain suppression and touch reward
- Oxytocin differentially modulates excitatory and inhibitory spinal neurons to rebalance ascending pain and touch pathways

## Introduction

Social bonds confer survival advantages across mammalian species, from enhanced protection against predators to improved resource acquisition and cooperative behaviors1–4. Touch serves as a primary sensory channel through which these bonds are formed and maintained, with mounting evidence that social touch ameliorates both emotional distress and physical pain5–9. Kangaroo care reduces pain during medical procedures in premature infants, while gentle stroking modulates noxious-evoked brain activity in human infants and adults10–15. Yet a fundamental puzzle remains: how do internal states—whether triggered by rewarding social contact or by injury and pain—modulate somatosensory processing to promote survival-enhancing social behaviors?

The neuropeptide oxytocin (OT) has emerged as a key orchestrator of prosocial behaviors, with hypothalamic OT neurons responding to social stimuli and projecting throughout the neuraxis6,16,17. Intriguingly, OT systems are activated not only during positive social interactions but also during pain and distress, suggesting a dual function18,19. While intrathecal OT shows promise for treating chronic pain in humans20, and preclinical studies demonstrate analgesic effects in acute pain models 21–24, the circuits underlying these effects—and critically, whether OT also modulates the rewarding aspects of social touch—remain poorly understood. This raises a central question: could OT function as a state-dependent neuromodulator that biases sensory processing toward prosocial outcomes regardless of whether the triggering experience is pleasant or aversive?

Emerging evidence reveals the spinal dorsal horn as an active site where contextual signals modulate sensory processing. Placebo analgesia, cognitive expectations, and even social context (self-versus other-touch) measurably alter dorsal horn responses, demonstrating integration of internal states with incoming sensory information25–28. Yet the circuit mechanisms underlying this integration remain poorly understood. The spinal dorsal horn—the first site of central somatosensory integration—contains molecularly distinct projection neurons that form parallel ascending pathways29–31. These pathways encode opposing valences: GPR83+ spinoparabrachial neurons promote positive reinforcement while TACR1+ neurons drive avoidance behavior31. However, how descending neuromodulatory systems—particularly those governing internal state changes—influence these valence-encoding circuits to integrate motivational context with sensory processing remains unknown. Moreover, while peripheral C-low threshold mechanoreceptors (C-LTMRs) mediating pleasant touch32–49 and supraspinal OT systems regulating social homeostasis have been well characterized6,8,17,50,51, the spinal mechanisms linking these systems have not been identified.

Here we demonstrate that spinal oxytocin circuits serve as a fundamental prosocial mechanism that transforms tactile experiences into opportunities for enhanced social connection. Using mouse genetic tools, behavioral assays, calcium imaging, chemo- and optogenetics, slice electrophysiology, and human psychophysical studies, we show that both rewarding touch and noxious stimuli activate hypothalamic OT neurons projecting to the spinal dorsal horn, where they differentially modulate distinct valence-associated pathways. Spinal OT selectively enhances GPR83+ spinoparabrachial neuron responses to pleasant touch while inhibiting TACR1+ neuron responses to noxious inputs—simultaneously making social touch more rewarding and pain less aversive. Critically, spinal OTR expression is necessary for gentle touch to induce reward, establishing that affective touch processing occurs at the earliest stage of central integration. We find that mouse and human spinal OTR neurons maintain remarkably similar excitatory-inhibitory balance despite 90 million years of divergence, and that massage-induced OT release in humans correlates with reduced stimulus aversiveness and slowed spinal somatosensory processing. These findings reveal an evolutionarily conserved feedback loop where affectively significant tactile experiences—whether positive or negative—trigger OT release that primes spinal circuits to enhance subsequent social engagement, identifying novel therapeutic targets for chronic pain and sensory processing disorders.

## Results

### Somatosensation across the valence spectrum correlates with OT release

Previous studies have shown that OT is released during positive social interactions, including tactile stimulation. We therefore designed a controlled study to determine if massage could induce OT release and allow investigation of rewarding touch effects on human somatosensory processing. Young females participated in two counterbalanced sessions consisting of either 30-minute skin-to-skin light pressure back massage or control questionnaire completion session (Figure 1a).

**Figure 1:**
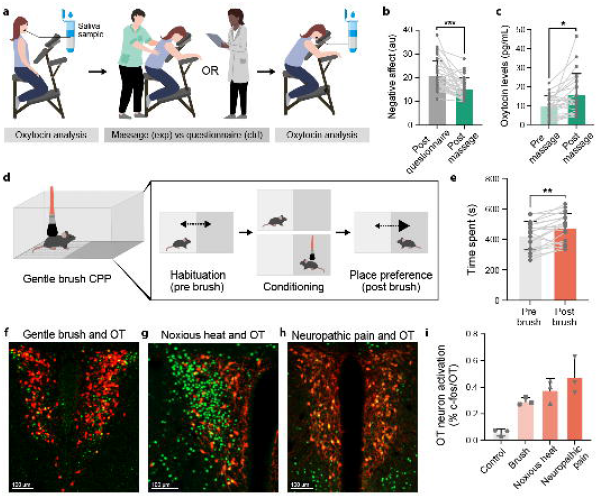
Somatosensation across the valence spectrum correlates with OT release. a. 37 young females participated in two counterbalanced sessions consisting of either 30-minute skin-to-skin light pressure back massage or control questionnaire completion. Salivary samples were collected before and after massage and questionnaire. Participants completed the positive and negative affect schedule questionnaire to assess their negative affective state during each session. b. Negative affect (au) was reported as significantly (t(29) = 4.339, p < 0.001) reduced during the massage session (mean = 14.65) compared to the control session (mean = 20.23). c. A significant increase in salivary oxytocin was observed after massage (mean = 15.84 pg/mL) compared to before massage (mean = 9.76 pg/mL; Tukey FWE-corrected post hoc comparison, p = 0.007). d. Schematic showing the gentle brush CPP paradigm: mice baseline preference was assessed over three days, then gentle brush conditioning was performed in the least preferred chamber for 12 days, with no stimulation in the preferred chamber, followed by a post-conditioning preference test. e. Mice developed gentle brush induced place preference in uninjured condition. Mean time spent pre-brush in least preferred chamber at baseline = 423.96 s, mean time spent post-brush in least preferred chamber at baseline = 473.26 s, n=10 mice. Slice immunohistochemistry on OT-Cre; LSL-Tomato brain tissue using c-fos (green) and tomato (red) staining shows gentle brush (f), noxious heat (g) and neuropathic pain (h) correlate with activation of OT+ neurons in the paraventricular nucleus of the hypothalamus. i. Quantification of c-fos+ OT+ amongst total OT+ neurons shows a significant increase in activation of OT+ neurons by gentle touch (mean = 29.3%), noxious heat (mean = 37.69%) and neuropathic pain (mean = 47.53%) compared to control condition (mean = 5.74%). n=3 mice per condition. Error bars indicate standard deviation.

We hypothesized that a pleasant massage would increase OT levels. We collected saliva samples to measure OT52 and cortisol levels before and after each intervention, along with heart rate measurements and assessment of negative affect and state of relaxation to assess the effect of massage on internal state. Massage significantly improved the participants’ mood with a significant reduction in negative affect (Fig.1b), increased relaxation (Fig.S1a) and increased salivary OT levels immediately (Fig.1c) and one hour post-treatment compared to baseline and control conditions (Fig.S1b), while reducing cortisol levels (Fig.S1c) and heart rate (Fig.S1d). All participants rated the back massage as pleasant (Fig.S1e), confirming that we can induce OT release via pleasant massage in a controlled setting.

These results establish that positively valenced tactile stimulation correlates with OT release in humans. To test whether this correlation exists in mice, we first used social touch-like stimulation and quantified its rewarding properties using conditioned place preference (CPP) (Fig.1d). As previously shown53–55, gentle brushing of back hairy skin during successive conditioning sessions induced significant place preference (Fig.1e), indicating that mice find social touch-like stimulation rewarding. Importantly, this reward persisted in chronic neuropathic pain conditions (irreversible Spared Nerve Injury neuropathic pain model, Fig.S1f), similar to the known benefits of comforting touch for humans experiencing pain.

OT is released centrally by neurons in both the paraventricular nucleus (PVN) and supraoptic nucleus (SON) of the hypothalamus. Previous studies have shown that PVN OT neurons are activated by various social behaviors. We therefore examined whether rewarding touch activates PVN OT neurons using OT-Cre mice crossed with cre-dependent tomato reporter mice. Rewarding touch activated 29% of PVN OT neurons compared to 6% at baseline (Fig.1f,i). Interestingly, similar to the dual role of OT in both maternal behavior and aggression in social behavior, we found that noxious heat stimulation (55°C) and post-spared nerve injury mechanical stimulation of hypersensitive hindpaws also robustly activated PVN OT neurons (37% and 47% activation respectively, Fig.1g,h,i). These findings demonstrate that changes in internal state induced by either positive or negative valence tactile experiences trigger OT release.

### Descending projections of PVN OT neurons overlap with OTR-expressing interneurons in the superficial dorsal horn and are activated by both rewarding and noxious tactile stimuli

The neural mechanisms underlying touch-induced analgesia remain debated, with some studies finding unchanged cortical pain representations despite reduced pain perception56, while others demonstrate cortical modulation57. These discrepancies suggest that early sensory processing sites, particularly the spinal dorsal horn, may play a critical role in touch-pain interactions. Recent evidence demonstrates that cognitive states modulate spinal processing. Placebo analgesia and prior knowledge about stimulus timing measurably alter dorsal horn responses25,26. Remarkably, self-touch versus other-touch shows different processing speeds already at the cervical spinal level, with faster somatosensory-evoked potentials for social touch28. These findings establish the spinal cord as an active site where internal states shape early sensory processing.

Altogether, these studies led us to hypothesize that the dorsal horn might serve as an active computational hub where oxytocin-mediated internal state changes could dynamically reconfigure how tactile inputs are processed before they reach the brain. After validating OT release during pleasant touch in humans, we found that both rewarding and noxious stimuli - despite their opposing valence - activate OT neurons in mice, suggesting the OT system responds to internal state changes regardless of stimulus quality. We next wanted to investigate whether this might translate into OT-mediated modulation of somatosensation at the earliest stage of tactile processing. First, we looked at the receptors for OT in the spinal cord. OT has 4 main receptors: oxytocin receptors (OTR), as well as vasopressin receptors 1a (AVPR1A), 1b (AVPR1B) and 2 (AVPR2). According to previous studies, the selectivity of OT for OTRs appears to be 10-100 times higher than for AVPRs and RNAsequencing data on mouse and human spinal cord tissue shows that spinal expression levels of AVPR1B, AVPR2 are too low to be detected58,59, and AVPR1A is expressed at very low levels which we confirmed using RNAscope (Fig.S2a). For these reasons, we decided to focus on OTR expression in the spinal cord. To do this we used RNA expression studies across species (mouse, rat, human) to establish the anatomy of spinal OTR circuits.

We hypothesized that OTRs would be strategically positioned in spinal regions that process both touch and pain information. It is generally considered that superficial layers of the dorsal horn of the spinal cord process and encode noxious mechanical, thermal and pruriceptive stimuli, while intermediate and deeper layers process light touch and proprioceptive information. Using RNAscope on postmortem human tissue, we found OTR mRNAs broadly present in the dorsal horn of both male and female human spinal cord (Fig.2a). 29% of dorsal horn neurons express OTRs and 3% are high expressors predominantly located in superficial and intermediate layers (Fig.2b). RNAscope on rat and mouse spinal cord tissue confirmed OTR expression patterns in superficial dorsal horn layers (Fig.S2b,c).

**Figure 2:**
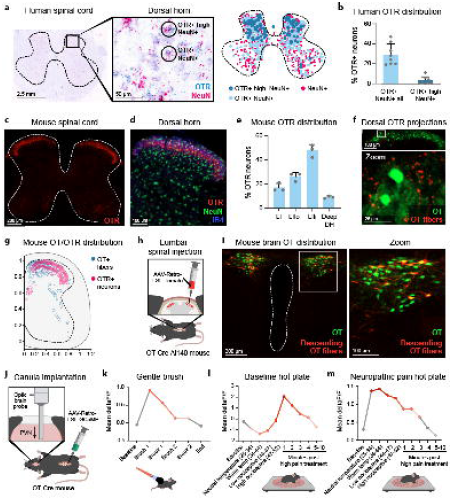
Descending projections of PVN OT neurons overlap with OTR-expressing interneurons in the superficial dorsal horn and are activated by both rewarding and noxious tactile stimuli. a. Chromogenic in situ hybridization (RNAscope) was performed on human lower thoracic to lumbar spinal cord tissue harvested from brain-dead organ-donors (50-80 years old, 2 males 1 female). OTR and NeuN probes were used and revealed a broad pattern of OTR expression among spinal neurons, with a more precise pattern of high OTR expression levels in the superficial and deep dorsal horn. b. Quantification of OTR+ neurons among total neurons shows 29.12% of human spinal neurons express OTR at low and high levels, with 3% of high expressors. c. Slice immunohistochemistry on mouse OTR-tomato spinal cord tissue using tomato (red) antibody shows expression of OTR in the superficial dorsal horn in mice, with very low levels of tomato expression in the deep dorsal horn and the ventral horn. d. Slice immunohistochemistry on mouse OTR-tomato dorsal horn spinal cord tissue using tomato (red), NeuN (green) antibodies and Isolectin Binding 4 (blue) to define lamina II outer shows that OTR is expressed in superficial dorsal horn neurons above, in and below lamina II outer. e. Quantification of OTR+ NeuN+ neurons per lamina of the dorsal horn amongst total OTR+ neurons shows that 17% of OTR neurons are located in lamina I, 26% in lamina II outer, 47.67% in lamina II inner and 9% in the deep dorsal horn. f. Slice immunohistochemistry on mouse OT-Cre; LSL-tomato; OTR-Venus dorsal horn spinal cord tissue shows projections from OT+ neurons to the superficial dorsal horn where OTR+ neurons are located. g. Density plots based on immunohistochemistry on OT-Cre; LSL-tomato; OTR-Venus spinal cord tissue show considerable overlap between OT+ fibers (blue) and OTR+ neurons (red) in the superficial dorsal horn. h. Schematic of mouse superficial lumbar spinal injections with AAV-Retro-LSL-Cherry in OT-Cre; Ai140 (LSL-GFP) mice to label spinal-projecting OT-expressing neurons. i. Slice immunohistochemistry on mouse OT-Cre; Ai140 brain tissue following superficial lumbar spinal injections with AAV-Retro-LSL-Cherry shows that a subset of hypothalamic paraventricular nucleus OT+ neurons (green) project to the dorsal horn of the spinal cord (red). The third ventricle is outlined in white dashed lines. j. Schematic showing OT-Cre mouse superficial lumbar spinal injection with AAV-Retro-GCaMP followed by paraventricular nucleus optic probe implantation to express GCaMP in spinal-projecting OT+ neurons and record their activity during various innocuous and noxious somatosensory stimuli. k. Variation of spinal-projecting OT+ neurons mean GCaMP fluorescence during gentle brush of back hairy skin showing activation of spinal projecting OT-expressing neurons during gentle brushing. l. Variation of spinal-projecting OT+ neurons mean GCaMP fluorescence during the dynamic hot plate assay (heat ramp ranging from room temperature to 52 degrees Celsius) in uninjured condition showing activation of spinal projecting OT-expressing neurons during noxious heat stimulation. m. Variation of spinal-projecting OT+ neurons mean GCaMP fluorescence during the dynamic hot plate assay (heat ramp ranging from room temperature to 52 degrees Celsius) in neuropathic pain condition (Spared Nerve Injury) showing expanded activation of spinal projecting OT-expressing neurons during innocuous and noxious heat stimulation in mice suffering from neuropathic pain. Error bars indicate standard deviation.

**Figure 3:**
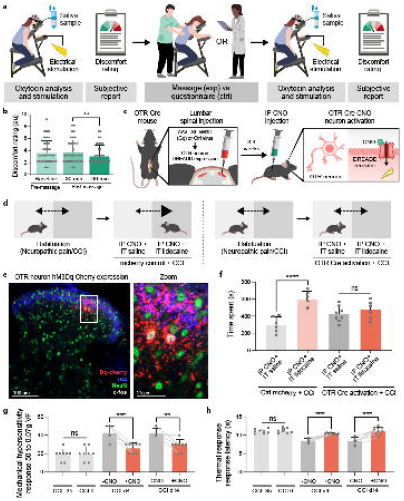
Spinal OTR neuron activation alleviates affective and sensory aspects of chronic neuropathic pain. a. 37 young females participated in two counterbalanced sessions consisting of either 30-minute skin-to-skin light pressure back massage or control questionnaire completion, coupled with electrical skin stimulation on the inside of the left forearm (wrist level), targeting the ulnaris nerve, before and after massage or questionnaire completion. Participants completed subjective reports rating the level of discomfort induced by the electrical skin stimulation before and after massage or questionnaire completion. b. Subjective discomfort ratings changed across time, with a reduction at later timepoints within the massage condition (Tukey FWE-corrected, p = 0.010 between early and later time-points). c. Schematic showing OTR-Cre mouse superficial lumbar spinal injection with AAV-LSL-hM3Dq-Cherry or control AAV-LSL-Cherry to induce Cre-dependent excitatory DREADD-Cherry or simple Cherry expression in superficial spinal OTR+ neurons, which will allow OTR neuron activation following clozapine N oxide (CNO) intraperitoneal (IP) injection. d. Schematic of a modified CPP assay to test the analgesic potential of superficial OTR neuron DREADD activation during neuropathic pain (Chronic Constriction Injury of the sciatic nerve). Mice are habituated to the 2 chamber setup and their baseline preference is measured. During the conditioning phase, mice receive either IP CNO and intrathecal saline injections in one chamber or IP CNO intrathecal lidocaine in the other chamber during counterbalanced morning and afternoon sessions over two days. Chamber preference post-conditioning is then measured again. e. Slice immunohistochemistry using Cherry (red), NeuN (green), and c-fos (white) antibodies and IB4 staining (blue) on mouse OTR-Cre spinal dorsal horn tissue following superficial lumbar spinal injections with AAV-LSL-hM3Dq-Cherry and, weeks later, IP CNO injection, shows expression of the hM3Dq (excitatory DREADD) virus in superficial spinal OTR neurons and activation of OTR+ cherry+ neurons located in lamina II of the superficial dorsal horn by CNO, with lamina II outer delimited by IB4 positive staining. f. Quantification of time spent by control and experimental mice suffering from neuropathic pain in chambers associated with either activation of superficial spinal OTR neurons only (IP CNO + IT saline) or activation of superficial spinal OTR neurons + lidocaine induced pain reduction (IP CNO + IT lidocaine). Quantification shows that experimental mice show no preference for activation of superficial spinal OTR neurons + lidocaine induced pain reduction over activation of superficial spinal OTR neurons only, contrary to control mice, suggesting that activation of superficial spinal OTR neurons has an analgesic effect comparable to intrathecal lidocaine on affective-motivational aspects of neuropathic pain. g. Quantification of mechanical sensitivity during 0.07g Von Frey filament stimulation of experimental mice in uninjured and neuropathic pain (CCI) conditions before and after intraperitoneal CNO injection. Quantification shows that the percentage of positive responses to the filament is increased due to neuropathic pain showing well described mechanical allodynia. This is reduced following CNO injection suggesting that OTR neuron activation can alleviate neuropathic pain-induced sensory-reflexive mechanical allodynia. h. Quantification of thermal sensitivity during the noxious heat Hargreaves assay on experimental mice in uninjured and neuropathic pain (CCI) conditions before and after intraperitoneal CNO injection. Quantification shows that the thermal response latency is decreased due to neuropathic pain showing well described thermal hyperalgesia. This is reduced following CNO injection suggesting that OTR neuron activation can alleviate neuropathic pain-induced sensory-reflexive thermal hypersensitivity. Error bars indicate standard deviation.

In mice, dorsal horn neurons show distinct functional specializations. Lamina I neurons respond to noxious mechanical, thermal and pruriceptive stimuli, while lamina II outer neurons encode both noxious and massage-like inputs35,60–66. Lamina II inner neurons receive socially salient tactile inputs and gate noxious signals, whereas deeper dorsal horn laminae process more discriminative touch and proprioceptive information. Using newly generated OTR-tomato mice and established OTR reporter lines, we found that OTR expression patterns in the superficial dorsal horn are consistent across cervical, thoracic and lumbar spinal levels in male and female mice (Fig.2c, Fig.S2d-h). We then characterized the dorsal spinal OTR neuron population and found that 92% of OTR-expressing spinal neurons reside in the two most superficial dorsal horn layers. 18% of OTR neurons were located in lamina I, 26% in lamina II outer, and 48% in lamina II inner (Fig.2d,e). Moreover, OTR neurons represented 17% of lamina I neurons, 43% of lamina II outer neurons and 53% of lamina II inner neurons (Fig.S2h).

The superficial dorsal horn in both mouse and human contains diverse interneuron populations that process touch and pain information. Using immunohistochemistry and RNAscope, we found that superficial OTR neurons overlap with numerous interneuron markers associated with pain and touch processing: calretinin (52%), SST (35%), calbindin (20%), PKCgamma (16%), GRPR (14%), GRP (9%) as well as minor overlap of 2% or less with parvalbumin, NK1R, neuropeptide Y (NPY), mu opioid receptor (MOR), delta opioid receptor (DOR), kappa opioid receptor (KOR) (Fig.S2.i-s)

To map descending OT fibers, we used OT-Cre mice crossed with cre-dependent synaptophysin-tomato reporter mice to label descending hypothalamic OT neurons synapses, combined with OTR-venus reporter mice to label spinal OTR neurons. Consistent with previous findings in rats, we found that PVN OT neurons project to the superficial dorsal horn and we identified a substantial overlap with superficial OTR expressing neurons (Fig.2f,g). Retrograde viral tracing using spinal injection of a retrograde Cre-dependent tomato virus in OT-Cre; Cre-dependent GFP reporter mice confirmed that a subset of PVN OT neurons projects to the spinal cord (Fig.2h,i). We then used retrograde Cre-dependent GCaMP expression in OT-Cre mice with fiber photometry to identify the response of these descending OT fibers to tactile stimuli along the valence spectrum (Fig.2j). Recordings revealed strong activation of descending OT neurons by rewarding touch, the social touch-like stimulus used in our CPP assay (Fig.2k) as well as noxious heat (47-52°C) at baseline, with activation persisting beyond stimulus offset (Fig.2l). Interestingly, the activation profile of descending OT neurons is expanded in neuropathic pain conditions with stronger and broader responses to both innocuous and noxious temperature after SNI (Fig.2m). These anatomical and functional findings position spinal OT-OTR circuits to modulate both pain and affective touch experiences at the earliest stage of somatosensory processing.

### Spinal OTR neuron activation alleviates affective and sensory aspects of chronic neuropathic pain

Our findings demonstrate that both rewarding touch and noxious stimuli activate PVN OT neurons which project to the superficial dorsal horn. Moreover, this region of the spinal cord is enriched with OTR-expressing interneurons that process pain and touch information. This anatomical convergence suggests that OT release in the spinal cord could function to scale a range of tactile experiences from pleasant to unpleasant, for example reducing the aversive impact of painful experiences. Pain reduction operates through distinct mechanisms: decreased signal intensity (sensory-discriminative) or reduced negative valence (affective-motivational)67. At the peripheral level, nociceptor sensitization and ion channel function determine signal strength60. Spinal circuits can then gate sensory transmission, as demonstrated by placebo-induced BOLD signal reduction in dorsal horn neurons25. From the spinal cord to the brain, parallel ascending pathways diverge: spinothalamic projections to somatosensory cortex encode intensity, while spinoparabrachial-amygdala circuits correlate with affect, though new studies suggest spinothalamic projections also play a role in valence encoding29,31. Critically, basolateral amygdala ensembles specifically encode pain unpleasantness—their silencing abolishes aversion without altering reflexive withdrawal68. CGRP neurons in thalamic SPFp exemplify this distinction, inducing aversive memory without changing mechanical thresholds29,68. Thus pain comprises dissociable components—intensity and negative valence—processed through distinct but overlapping spinal circuits. This anatomical organization—with both dimensions represented at the earliest stage of central processing—makes the spinal oxytocin system uniquely positioned to modulate both how much something hurts and how unpleasant it feels.

First we used our human massage study to assess whether massage-induced OT release could be linked to reduced negative valence of aversive somatosensory stimuli. To do this, we included skin electrical stimulation of the ulnar nerve at the level of the wrist and asked participants to provide subjective ratings of unpleasantness and pain during the control and massage sessions (Fig.3a). Participants reported the electrical stimulation as uncomfortable though not painful. Interestingly, we found that participants overall rated the stimulation as significantly less uncomfortable after massage and OT release (Fig.3b) but not after filling out questionnaires during the control session (Fig.S3a, t2 electrical stimulation immediately after massage vs t3 electrical stimulation 1h after massage, mean difference = 0.6, ptukey= 0.01 Post Hoc Comparisons - Treatment ✻ SEP subjective Time). This suggests that massage-induced OT release was associated with reduced negative valence of an aversive tactile stimulus.

Clinical trials show promise for intrathecal OT injections in treating chronic neuropathic pain20,69 and preclinical studies show analgesic benefits of spinal OT-OTR circuit activation in acute pain conditions19,21–24, pointing to underlying spinal mechanisms of OT mediated analgesia. To understand these mechanisms, we first used patch-clamp electrophysiology to assess the overall effect of OT release on OTR neurons. We found that OT depolarizes OTR spinal neurons (7/12 OTR+ neurons), increasing their action potential firing probability and frequency (Fig.S3b). To test whether this increased excitation translates to behaviorally relevant analgesia, we then employed excitatory DREADD (Designer Receptors Exclusively Activated by Designer Drug) technology to mimic this effect by injecting cre-dependent hM3Dq-cherry (OTR-Dq) or control cherry virus into the superficial dorsal horn of OTR-Cre mice (Fig.3c, Fig.S3c). Inducing neuropathic pain (chronic constriction injury (CCI)), we tested whether spinal OTR circuit activation could improve multiple dimensions of the pain experience.

Pain encompasses both affective-motivational and sensory-discriminative components that can be independently assessed in rodents. We first examined the affective-motivational dimension using conditioned place preference at 14 days post-CCI during ongoing neuropathic pain. We used intraperitoneal injection of the DREADD receptor synthetic ligand Clozapine-N-oxide (CNO), to test the analgesic potential of OTR neuron activation, coupled with either intrathecal saline injection or intrathecal lidocaine injection as a positive control for pain relief (Fig.3d,e). Control mice, which did not express the excitatory DREADD receptor Dq, showed strong preference for chambers paired with CNO-lidocaine (which relieves ongoing pain) over CNO-saline. However, OTR-Dq mice showed no chamber preference, indicating that spinal OTR activation reduced the negative valence of neuropathic pain to levels comparable to lidocaine treatment (Fig.3f).

We next assessed the analgesic potential of OTR neuron activation on the sensory-discriminative dimension of ongoing neuropathic pain using mechanical and thermal hypersensitivity assays. CNO injection in OTR-Dq mice significantly reduced both mechanical and thermal hypersensitivity at acute (4 days) and chronic (14 days) timepoints post-CCI compared to controls (Fig.3g,h). These analgesic effects are consistent with the known efficacy of OT in acute pain models and demonstrate that spinal OTR activation can modulate pain intensity as well as pain affect.

These results demonstrate that spinal OT release—which can be triggered by either rewarding touch or noxious experiences—can modulate both affective-motivational and sensory-discriminative aspects of pain. This data together with our findings of reduced negative valence of aversive stimuli in humans supports our hypothesis that the spinal cord could serve as a key site where OT modulates both affective-motivational and sensory-discriminative dimensions of chronic neuropathic pain.

### Spinal OTR expression is necessary for rewarding aspects of gentle touch

The rewarding nature of gentle touch is fundamental to neural and social development, physical health, and mental well-being, as highlighted during recent social isolation experiences. While the skin and brain circuits underlying gentle touch have been conserved across evolution40,42,48,49,70–73, a fundamental question remains: do spinal processing centers actively contribute to the rewarding quality of gentle touch, or do they merely relay sensory information to the brain for affective processing? In particular, given the long-standing bias toward brain-based neuromodulation of sensory processing, the potential role of descending neuromodulation in shaping how the spinal cord processes rewarding touch has remained largely unexplored. Recent evidence demonstrates that the dorsal horn actively processes contextually distinct types of touch, with self-produced and other-produced touch being differentiated already at the cervical spinal level through faster somatosensory-evoked potential latencies for social touch compared to self-touch28. This suggests that spinal circuits coupled with descending modulation can distinguish between touch contexts that carry different social and affective significance.

Building on our evidence that both rewarding and noxious tactile stimuli activate PVN OT neurons projecting to OTR-expressing superficial dorsal horn circuits, we found that massage increases OT release and reduces discomfort ratings of aversive stimuli in humans while it reduces both sensory and affective responses to aversive stimuli in mice. Notably, 74% of spinal OTR neurons are located in lamina II, which receives C-low threshold mechanoreceptor (C-LTMR) inputs that mediate pleasant touch sensations32,33,35–39,48,49,74. This anatomical positioning suggests that spinal OT-OTR circuits could directly modulate the rewarding properties of gentle touch at the earliest stage of somatosensory processing. Our converging evidence—that massage increases OT release, that OTR neurons are strategically positioned in touch-processing laminae, and that spinal OTR activation reduces pain aversiveness—led us to hypothesize that descending OT modulates spinal processing of gentle touch in ways that influence the overall positive valence ultimately assigned to the experience.

Using the social touch-like CPP assay established earlier, we tested whether spinal OTR signaling is necessary for touch reward. We used a conditional knockdown approach, injecting either Cre-expressing (OTR cKO) or control Cherry-expressing viruses into the superficial dorsal horn of OTR-flox mice at the lower thoracic-upper lumbar level, which receives inputs from back hairy skin where the brush was applied (Fig.4a). RNAscope validation confirmed that knockdown was restricted to the injection site (Fig.4b,c), with no changes in OTR expression at cervical levels (Fig.S4a), ensuring that supraspinal OT system function remained intact.

**Figure 4:**
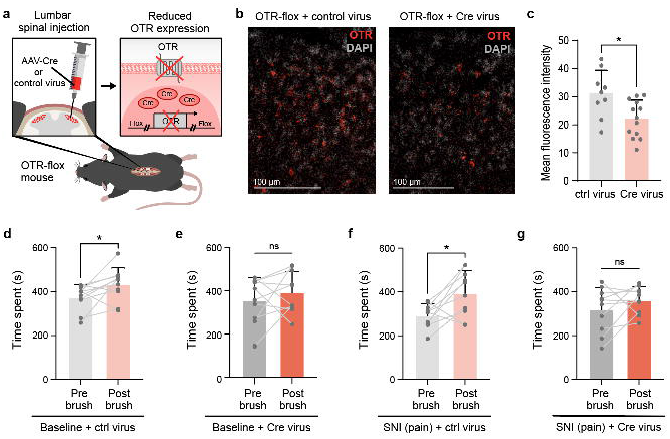
Spinal OTR expression is necessary for rewarding aspects of gentle touch. a. Schematic showing OTR-flox mouse superficial lower thoracic to upper lumbar spinal injections with AAV-Cre or control virus to induce localised knock down of OTR expression in superficial spinal OTR+ neurons. b. RNAscope using a mouse OTR probe (red) on OTR RNA expression in OTR-flox + Cre virus mouse superficial dorsal horn slice tissue compared to OTR-flox + control virus mouse superficial dorsal horn slice tissue, counterstained with nucleus binding DAPI (light grey). c. Quantification of mean fluorescence intensity of OTR-flox + Cre virus mouse superficial dorsal horn tissue compared to OTR-flox + control virus mouse superficial dorsal horn tissue using RNAscope with a mouse OTR probe shows reduced OTR expression in Cre virus condition compared to control virus. d-g. Quantification of time spent by control and OTR spinal knockdown mice in the gentle brush associated chamber pre- and post-conditioning using the gentle-brush CPP assay described in Figure 1. d. Uninjured control mice show gentle brush-induced place preference. e. Uninjured spinal OTR knockdown mice do not show gentle brush-induced place preference. f. Control mice suffering from neuropathic pain (Spared Nerve Injury) show gentle brush-induced place preference. g. Spinal OTR knockdown mice suffering from neuropathic pain (Spared Nerve Injury) do not show gentle brush-induced place preference. This suggests that correct levels of OTR expression in superficial spinal OTR neurons is required for gentle brush-induced place preference and gentle touch reward. Error bars indicate standard deviation.

Gentle brushing of back hairy skin induced robust reversal of place preference in control mice both at baseline and after spared nerve injury (Fig.4d,f), a mouse model of irreversible neuropathic pain, confirming the persistent rewarding nature of social touch even during chronic pain75–78. Conversely, spinal OTR knockdown abolished gentle brush-induced place preference under both conditions (Fig.4e,g), despite preserved tactile sensitivity as confirmed by back hairy skin tactile prepulse inhibition assays (Fig.S4b). This demonstrates that spinal OTR circuits specifically contribute to the affective-motivational aspects of touch rather than its basic sensory detection.

These findings establish that OT signaling specifically in the spinal cord is essential for the rewarding aspects of gentle touch at baseline and its therapeutic benefits in chronic pain conditions. This regional specificity of OTR function is consistent with findings in other CNS regions, where local OTR signaling mediates distinct behavioral outcomes: hippocampal OTR knockdown impairs social but not general memory formation79, while ventral tegmental area OTR activation specifically enhances social reward without affecting non-social motivation80 and nucleus accumbens OTR signaling gates critical periods for social reward learning81. Our results extend this principle to the spinal cord, demonstrating that descending OT modulation at the first central relay can shape affective-motivational processing traditionally thought to require mainly supraspinal evaluation. This aligns with emerging evidence that spinal circuits actively contribute to complex behaviors: spinal responses correlate with placebo analgesia25,82 and cognitive expectation exerts top-down control over spinal sensory processing to shape perception26. Rather than functioning as a passive relay, the spinal cord appears to serve as an active computational hub where descending neuromodulators like oxytocin can fundamentally reshape how sensory experiences acquire motivational and affective significance.

### Spinal OT balances inhibition and excitation in dorsal horn circuits

Having established that spinal OT release both reinforces rewarding touch and reduces the negative valence of aversive and painful tactile stimuli, we sought to elucidate the neural circuitry underlying this functional dichotomy. Internal states—such as social isolation, exclusion, and pain— modulate excitation/inhibition (E/I) balance in neural circuits across multiple brain regions83. For example, newly identified social isolation-activated neurons in the medial preoptic nucleus are glutamatergic while most social reunion-activated neurons are GABAergic. Interestingly, isolation also leads to activation of OT neurons in the PVN and administration of an oxytocin receptor antagonist during isolation reduces social rebound during reunion6. Oxytocin signaling itself can provide a key mechanism for E/I modulation: in the auditory cortex, oxytocin receptor activation rebalances cortical E/I dynamics by enhancing recruitment of inhibitory interneurons to improve signal-to-noise ratio, thereby enabling complex behaviors such as maternal responses to pup calls17. Meanwhile, in the insular cortex, oxytocin signaling increases during social exclusion and acts protectively against subsequent physical pain, demonstrating how neuromodulators can bridge different temporal scales—modifying neural processing to link social experiences with later sensory perception18. However, whether internal state changes mediated by oxytocin can modulate E/I balance in spinal cord somatosensory circuits remains unexplored. Recent work shows that expectation influences initial sensory processing in the human spinal cord, linking cognitive state changes with top-down control of dorsal horn sensory processing26. We hypothesized that OT achieves its dual functions—enhancing rewarding touch while reducing pain—by differentially modulating excitatory and inhibitory spinal circuits in the dorsal horn.

Using RNAscope on mouse and rat tissue, we found that spinal OTR neurons are composed of 58% excitatory (vGluT2+) and 38% inhibitory (VGAT+) populations in mice (Fig.5a,b,c) and 69% excitatory and 33% inhibitory populations in rats (Fig.S5a,b,c). Using the inhibitory neuron marker pax2 on spinal cord tissue from OTR-Venus and OTR-Tomato reporter mice also showed that 34% and 43% of superficial OTR neurons respectively have an inhibitory neurotransmitter profile based on histology studies (Fig.S5d,e,f). To validate this phenotype at a functional level, we performed slice electrophysiology recordings from OTR neurons in OTR-tomato; GAD67-GFP mice in the absence or presence of OT. Patch clamp recording revealed that excitatory OTR neurons (eOTR neurons: OTR-Tomato+ Gad67GFP-) typically exhibit delayed firing (69%,Fig.5d) while inhibitory OTR neurons (iOTR neurons: OTR-Tomato+ Gad67GFP+) predominantly show tonic firing patterns (75%,Fig.5e). Bath application of OT increased excitability in most inhibitory OTR neurons (71%) while producing variable effects on excitatory neurons (Fig.5f,g, Fig.S5g). This approach showed a stronger OT modulation of firing patterns of inhibitory OTR neurons (66% of all OTR neurons with a change in firing pattern) compared to excitatory OTR neurons (33%) (Fig.5h). These histological and electrophysiological findings suggest a slight functional bias towards increased inhibition in the dorsal horn in response to OT release.

**Figure 5:**
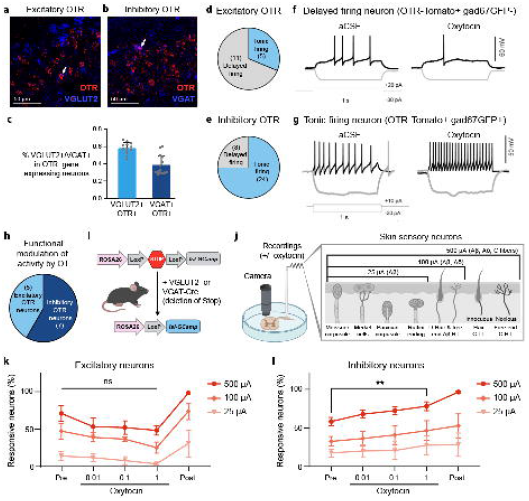
Spinal OT balances inhibition and excitation in dorsal horn circuits. a. RNAscope using a mouse OTR probe (red) and a mouse VGLUT2 probe on WT mouse superficial dorsal horn slice tissue shows overlap between OTR and VGLUT2 mRNA expression (white arrow). b. RNAscope using a mouse OTR probe (red) and a mouse VGAT probe on WT mouse superficial dorsal horn slice tissue shows overlap between OTR and VGAT mRNA expression (white arrow). c. Quantification of VGLUT2+ OTR+ neurons over total superficial spinal OTR+ neurons and VGAT+ OTR+ over total superficial spinal OTR+ neurons. d. Proportion of excitatory OTR-tomato+; gad67GFP-OTR neurons by firing pattern: the majority of excitatory superficial spinal OTR neurons show a delayed firing pattern (11) with only 5 showing a tonic firing pattern. e. Proportion of inhibitory OTR-tomato+; gad67GFP+ OTR neurons by firing pattern: the majority of inhibitory superficial spinal OTR neurons show a tonic firing pattern (24) with only 8 showing a delayed firing pattern. f. Representative traces of the modulatory effect of OT on an excitatory superficial spinal OTR neuron (OTR-tomato+; gad67GFP-) with a delayed firing pattern: OT reduces the firing rate. g. Representative traces of the modulatory effect of OT on an inhibitory superficial spinal OTR neuron (OTR-tomato+; gad67GFP+) with a tonic firing pattern: OT increases the firing rate. h. Quantification of the modulatory effect of OT on the firing rate of inhibitory OTR-tomato+; gad67GFP+ OTR neurons vs excitatory OTR-tomato+; gad67GFP-OTR neurons shows a bias towards OT modulating more inhibitory than excitatory OTR neurons. i. Schematic of the genetic strategy employed to assess OT modulatory effect on dorsal horn inhibitory and excitatory circuits: VGLUT2-Cre or VGAT-Cre mice were crossed to Cre-dependent GCaMP expressing mice to induce expression of the calcium sensor in excitatory or inhibitory neurons. j. Schematic of 2 photon calcium imaging of the response of excitatory or inhibitory dorsal horn circuits to dorsal root electrical stimulation in the absence or presence of bath oxytocin. Fresh superficial dorsal horn slices with dorsal roots still attached were stimulated at 25, 100 or 500 μA to recruit either Aβ fibers only, Aβ + Aδ fibers or Aβ+ Aδ + C fibers respectively. k. Percentage of superficial dorsal horn excitatory neurons responsive to 25, 100 and 500 μA dorsal root stimulation prior to and in the presence of 0.01, 0.1 and 1μM bath oxytocin, and following GABAergic and glycinergic spinal cord disinhibition with bath application of 10 μM bicuculline and 1 µM strychnine. l. Percentage of superficial dorsal horn inhibitory neurons responsive to 25, 100 and 500 μA dorsal root stimulation prior to and in the presence of 0.01, 0.1 and 1μM bath oxytocin, and following GABAergic and glycinergic spinal cord disinhibition with bath application of 10 μM bicuculline and 1 µM strychnine. Oxytocin significantly increases the response of superficial dorsal inhibitory circuits to 500 µA dorsal root stimulation. Error bars indicate standard deviation for panel c, and standard error of the mean for panels k and l.

To probe circuit-level effects, we used two-photon calcium imaging to record global superficial dorsal horn responses to graded dorsal root stimulation (25μA to recruit Aβ fibers, 100μA for Aβ+Aδ fibers, 500μA for all fibers including C-fibers) before and after OT bath application. We used acute lumbar spinal cord slice tissue of vGluT2-Cre and VGAT-Cre mice expressing the Cre-dependent fluorescent calcium sensor GCaMP6s to look at the effect of OT on excitatory vs inhibitory dorsal horn circuits (Fig.5i,j). We found an interesting differential effect of OT bath application with an overall trend toward reduced excitation and increased inhibition in response to sensory stimulation (Fig.5k,l). This modulation was statistically significant specifically during inhibitory responses to electrical stimulations which recruited all DRG fibers including C-fibers (500μA stimulation) (Fig.5l). This is notable because in mice C-fibers represent the majority of overall DRG inputs (over 80%) while Aβ and Aδ fibers only represent about 10% of DRG neurons respectively.

These findings reveal that OT rebalances dorsal horn circuits by enhancing inhibitory responses while suppressing excitatory responses, with preferential modulation of C-fiber inputs that could convey pleasant touch and/or nociceptive information. This circuit-level rebalancing provides a mechanism by which changes in internal state—triggered by either rewarding touch or noxious experiences—can modulate subsequent sensory processing at the earliest stages of somatosensory processing in the spinal cord.

### Spinal OT increases inhibitory neuron responses to noxious peripheral inputs to gate pain pathways while enhancing the recruitment of touch reward pathways

Our findings demonstrate that spinal OT acts as a versatile neuromodulator, orchestrating both the enhancement of rewarding touch and the attenuation of pain through differential recruitment of excitatory and inhibitory dorsal horn circuits in response to the activation of C-fibers. However, critical questions remain about the precise circuit mechanisms underlying these dual functions. Which specific C-fiber inputs are modulated by OT? How do inhibitory and excitatory OTR-expressing neurons integrate these different sensory modalities? And crucially, how does this excitatory/inhibitory balance translate into modulation of spinal output pathways to shape the experience of gentle touch and pain?

In mice, C-fibers represent 80% of somatosensory neurons and comprise a heterogeneous population responding to modalities ranging from aversive noxious heat (TRPV1+84,85) and mechanical stimulation (MrgD+86,87) to potentially pleasant light touch (TH+ C-LTMRs37,88) and massage (MrgB4+32,89). And OTR neurons are strategically positioned in spinal regions that process both light touch and noxious information. To understand which C-fiber inputs might be modulated by spinal OT release, we investigated the peripheral inputs to superficial OTR neurons. We used TRPV1-Cre, MrgD-CreER, MrgB4-Cre, and TH-CreER mouse lines coupled to synaptophysin reporters in OTR reporter mice to visualize their inputs onto OTR neurons (Fig.S6a). We found direct contacts from both light touch (MrgB4+ and TH+) and noxious (TrpV1+ and MrgD+) inputs onto OTR spinal neurons (Fig.S6b,c,d,e). We then used the same genetic approach to express channelrhodopsin in TrpV1+, MrgD+, MrgB4+ or TH+ fibers in OTR reporter mice combined with spinal cord slice electrophysiology to test the functional relevance of our anatomical findings (Fig.6a). We previously noted that a majority of excitatory OTRs (eOTRs) exhibit delayed firing patterns while inhibitory OTRs (iOTRs) have a tonic firing profile. We used these physiological characteristics to assess whether inputs conveying light touch or noxious stimuli show a connectivity bias towards putative eOTRs (delayed firing OTRs) or putative iOTRs (tonic firing OTRs). We found that OTRs are indeed synaptically coupled to TrpV1+, MrgD+, MrgB4+ and TH+ fibers because optogenetic activation of all four sensory neuron types evoked excitatory postsynaptic currents (EPSCs) in OTR neurons. Interestingly, both eOTR and iOTR populations received noxious and innocuous inputs with no obvious bias (Fig.6b).

**Figure 6:**
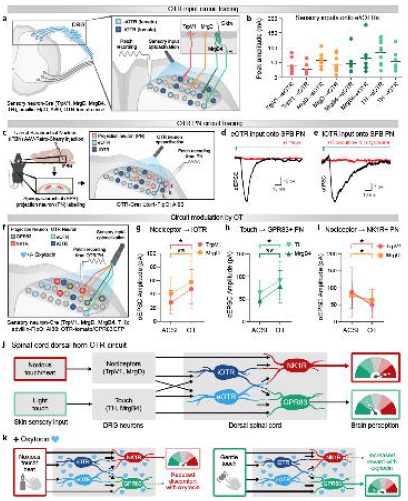
Spinal OT increases inhibitory neuron responses to noxious peripheral inputs to gate pain pathways while enhancing the recruitment of touch reward pathways. a. Schematic of slice electrophysiology patch clamp recording from OTR neurons coupled with optogenetic stimulation of sensory inputs. b. Quantification of the peak amplitude of opto-evoked sensory inputs onto putative excitatory and inhibitory OTR neurons based on their firing patterns. Both eOTRs and iOTRs receive TrpV+, MrgD+, MrgB4+ and TH+ sensory inputs. c. Schematic of slice electrophysiology patch clamp recording from retrogradely labelled spinoparabrachial (SPB) projection neurons (PNs) coupled with optogenetic stimulation of OTR neurons. d. Representative trace of opto-evoked eOTR inputs onto SPB PN. The excitatory synaptic transmission was blocked by bath application of the kainate/AMPA receptor antagonist CNQX. e. Representative trace of opto-evoked iOTR inputs onto SPB PN. The inhibitory synaptic transmission was blocked by bath application of the GABAA receptor/glycine receptor antagonists bicuculline/strychnine. f. Schematic of slice electrophysiology patch clamp recording from either OTR neurons classified based on their firing pattern or retrogradely labelled SPB PNs classified by their expression of the marker GPR83 coupled with optogenetic stimulation of sensory inputs. g. Quantification of the mean amplitude of TrpV1+ and MgrD+ opto-evoked post-synaptic currents in iOTR neurons before and after OT bath application shows increased amplitude in the presence of OT in response to noxious stimulus-sensitive sensory neuron inputs. h. Quantification of the mean amplitude of MrgB4+ and TH+ opto-evoked post-synaptic currents in GPR83+ SPB PNs before and after OT bath application shows increased amplitude in the presence of OT in response to gentle touch stimulus-sensitive sensory neuron inputs. i. Quantification of the mean amplitude of TrpV1+ and MgrD+ opto-evoked post-synaptic currents in GPR83-putative NK1R+ SPB PNs before and after OT bath application shows decreased amplitude in the presence of OT in response to noxious stimulus-sensitive sensory neuron inputs. j. Proposed architecture of the spinal dorsal horn OTR circuit inputs and outputs. k. Proposed circuit level mechanism of spinal OT release and its impact on conscious perception of noxious or gentle somatosensory inputs.

To understand how spinal OT might shape sensory processing, we wanted to identify the output of spinal OTR neurons. We used intersectional genetics: OTR-Cre with the dorsal horn-specific Lbx1-FlpO to express a Cre- and Flp-dependent synaptophysin-GFP in spinal OTR neurons and found that spinal OTR neurons terminate mostly in lamina I and lamina II outer (Fig.S6f). These superficial laminae contain anterolateral tract neurons that convey an array of thermal, mechanical and itch signals to supraspinal structures such as the thalamus and the parabrachial nucleus. The spinoparabrachial (SPB) pathway in particular contains ∼90% of spinal output neurons. To test whether spinal OTR neurons target SPB projection neurons, we used retrograde labeling from the lateral parabrachial nucleus (lPBN) to identify SPB projection neurons, combined with slice electrophysiology and using OTR-Cre x Lbx1FlpO mice to express a Cre- and Flp-dependent channelrhodopsin in spinal OTR neurons (Fig.6c). We found that optogenetic activation of spinal OTR neurons evoked excitatory and inhibitory post-synaptic currents in SPB neurons. We then used bath application of CNQX or bicuculline and strychnine to pharmacologically isolate excitatory and inhibitory connections, respectively. This showed that both eOTR and iOTR neuron subsets provide direct synaptic input to SPB projection neurons (Fig.6d,e).

Overall, we have established that both eOTR and iOTR spinal neurons receive a wide range of innocuous and noxious somatosensory inputs, and both eOTR and iOTR spinal neurons target projection neurons of the main superficial output pathway. This blueprint of OTR inputs and outputs reveals a complex substrate for how spinal OT might modulate both noxious and innocuous inputs to enhance touch reward and reduce pain. To test at a global level whether OT shapes the activation of output pathways by incoming somatosensory inputs, we used slice electrophysiology and intersectional optogenetics broadly targeting somatosensory neurons (cdx2-Cre x advillin-FlpO and Cre- and Flp-dependent channelrhodopsin) coupled with bath OT and retrograde labeling of SPB projection neurons (Fig.S6g). Recent work has identified that the activation of parallel spinoparabrachial (SPB) pathways correlates with distinct affective information: GPR83+ neurons can contribute to touch reward31,49 while NK1R+ neurons are associated with pain aversion30,31, together representing ∼90% of SPB neurons. We hypothesized that spinal OT shapes somatosensory processing at the circuit level by modulating how somatosensory inputs activate GPR83+ and/or NK1R+ SPB output pathways. To test this hypothesis, we used cdx2-Cre; advillin-FlpO; LSL-FSF-channelrhodopsin; GPR83GFP mice where somatosensory neurons broadly express channelrhodopsin and we can visualize GPR83+ neurons in green. Coupled with retrograde labeling of SPB neurons with retro AAV-cherry and bath OT application, we were able to record how OT modulates the response of GPR83+ SPB projection neurons (red and green) and putative NK1R+ SPB projection neurons (red only) to somatosensory input. While we did not observe a clear change in light-evoked ePSCs in GPR83+ SPB neurons, the response of putative NK1R+ SPB neurons to broad somatosensory stimulation was significantly dampened (Fig.S6h). This suggests an overall inhibitory effect of OT on NK1R+ output pathways, known to be associated with avoidance and nocifensive pain-associated behavior.

Following this, we wanted to tease apart the role of OT in modulating innocuous versus noxious somatosensory input. To test modality-specific OT effects, we performed a wide array of electrophysiology studies of the response of e/iOTR neurons to TrpV1/MrgD/MrgB4/TH input optogenetic activation and their potential modulation by bath OT (Fig.6f) using the OTR-tomato mouse line and OTR neurons firing patterns to record from e/iOTR neurons. Strikingly, OT increased the excitability of putative iOTR neurons (tonic firing) in response to both TRPV1+ and MrgD+ stimulation (Fig.6g) but had no effect on eOTR neuron responses to these noxious inputs (Fig.S6i). OT did not affect overall OTR neuron responses to light touch-sensitive TH+ or MrgB4+ stimulation (Fig.S6j,k). These results demonstrate that OT selectively enhances inhibitory circuit responses to noxious inputs, providing a mechanism for modality-specific modulation that reduces the impact of aversive stimuli.

We then examined how OT modulates specific output pathways by combining slice electrophysiology and optogenetic activation of C-fiber inputs (TRPV1+, MrgD+, MrgB4+, TH+) with recordings from genetically and retrogradely identified GPR83+ (red and green) and NK1R+ (red only) SPB neurons before and after OT application (Fig.6f). OT produced strikingly different effects on spinal outputs depending on input modality: it increased GPR83+ neuron responses to both massage-responsive MrgB4+ and gentle touch-responsive TH+ inputs (Fig.6h) while decreasing NK1R+ neuron responses to noxious heat-responsive TRPV1+ inputs (Fig.6i).

We found it intriguing that spinal OT release could lead to both increased inhibition of noxious inputs and increased activation of output pathways by gentle touch. The superficial laminae of the dorsal horn show a precise anatomical organization which correlates with different input modalities: more noxious mechanical, thermal and itch inputs in laminae I and IIo and more socially rewarding inputs like social touch in lamina II inner. To determine whether the neurotransmitter profile of OTR neurons is similar across laminae, we quantified the ratio of e/iOTR neurons using the classic marker of lamina II outer isolectin-B4 staining to identify lamina I (above IB4), lamina II outer (IB4+) and lamina II inner (immediately below IB4). Quantification using inhibitory marker pax2 revealed a dorsoventral gradient from predominantly iOTRs in lamina I (64.3% pax2+ OTR+ neurons/total OTR+ neurons, Fig.S6l) to a minority of iOTRs in lamina II inner (23.7% pax2+ OTR+ neurons/total OTR+ neurons, Fig.S6m). This anatomical organization suggests that spinal OT release differentially modulates noxious versus innocuous inputs through lamina-specific inhibitory-excitatory balance: increased inhibition of noxious processing in lamina I and enhanced excitatory signaling of pleasant touch in lamina IIi, consistent with our SPB output findings (Fig.6h,i).

Our findings reveal pathway-specific OT modulation: enhanced GPR83+ responses to pleasant touch paired with suppressed TACR1+ responses to noxious inputs (Fig.6j,k). This bidirectional modulation means that internal state changes—from either rewarding or aversive experiences—prime spinal circuits to amplify social touch reward while dampening pain aversion.

### Human spinal OTR circuits mirror mouse mechanisms

Having identified shared OTR dorsal horn circuits mediating touch reward and pain modulation in mice, we investigated whether similar mechanisms operate in humans. Using RNAscope on human spinal cord tissue (Fig.7a), we found that human spinal OTR neurons, like those in mice, are comprised of mixed excitatory (46% OTR+ vGluT2+ neurons) and inhibitory (54% OTR+ VGAT+ neurons) neurotransmitter phenotypes (Fig.7b). While expression was broader in human dorsal horn, high OTR expressors were most frequent in superficial layers, mirroring mouse and rat expression patterns.

We hypothesized that massage-induced OT release would modulate somatosensory processing specifically at the spinal level in humans, as suggested by our mouse circuit analysis. We recorded somatosensory evoked potentials (SEPs) at spinal (C2)28,90 and cortical levels in response to median nerve electrical stimulation before and after massage or questionnaire sessions (Fig.7c,d).

Post-massage, we observed significantly increased SEP signal latency at both spinal and cortical levels (Fig.7e, Fig.S7a). Post-questionnaire, only the SEP signal latency at the cortical level was increased (Fig.S7b, c). Critically, individual analysis revealed a positive correlation between OT release magnitude and increased SEP latency specifically at the spinal level (Fig.7f), but not at the cortical level. This correlation, combined with participants’ reduced discomfort ratings for electrical stimulation post-massage (Fig.3b), demonstrates a direct link between touch-induced OT release and spinal somatosensory modulation in humans.

These results highlight the conservation of OT-mediated somatosensory modulation mechanisms from mice to humans and establish spinal OT circuits as a fundamental mechanism linking rewarding touch to pain relief across species91.

## Discussion

Our findings reveal how changes in internal state—triggered by either rewarding touch or noxious experiences—modulate sensory processing through oxytocin-mediated rebalancing of spinal circuits. The key insight is that both affectively significant tactile experiences activate hypothalamic OT neurons, which then bias spinal processing toward prosocial outcomes by differentially modulating parallel ascending pathways: enhancing GPR83+ neuron responses to pleasant touch while suppressing TACR1+ neuron responses to noxious inputs. The net result is a circuit state that makes social touch more rewarding and pain less aversive, thereby promoting approach behaviors that facilitate social bonding and recovery.

### A complete circuit for social touch allostasis

While brain OT systems regulating social homeostasis and peripheral C-LTMR neurons detecting rewarding social touch have been well characterized, the spinal mechanisms linking these systems remained unknown. Our study provides the first identification of spinal OT circuits as essential mediators of social touch reward, closing a critical loop in social touch neurobiology. We demonstrate that spinal OTR expression is necessary for gentle touch to induce conditioned place preference, revealing that the affective quality of touch is actively shaped at the earliest stage of central processing rather than emerging solely from supraspinal evaluation. This complements recent work identifying spinal PROKR2+ neurons in touch reward signaling53 and extends our understanding by revealing descending OT modulation as a state-dependent mechanism that amplifies rewarding touch experiences.

Recent studies have established that spinal circuits actively integrate contextual information with sensory inputs. Placebo analgesia reduces spinal dorsal horn activity, and cognitive expectations modulate initial sensory processing at the spinal level. Our work identifies oxytocin as a specific molecular mechanism for this state-dependent modulation, showing how internal states triggered by tactile experiences translate into circuit-level changes. Based on our findings and recent literature, we propose a positive feedback loop: peripheral C-LTMR activation → spinal GPR83+ projection neurons → lateral parabrachial nucleus → (potentially via periaqueductal gray) → PVN OT neurons → descending spinal OT release → enhanced processing of subsequent rewarding touch. This circuit architecture provides a neurobiological basis for the clinical observation that gentle touch enhances the pleasantness of subsequent touch92.

### Novel mechanisms of spinal OT action: differential modulation of distinct valence-associated pathways

A major advance from our work is the demonstration that spinal OT differentially modulates recently-identified parallel spinoparabrachial pathways that can be associated with opposing valences. Choi and colleagues recently discovered that GPR83+ and TACR1+ spinal projection neurons project to distinct supraspinal targets through anatomically and functionally distinct circuits, with GPR83+ neuron activation associated with intensity-dependent behavioral outcomes (reward at low to moderate intensities, avoidance at high intensities) and TACR1+ neuron activation consistently associated with avoidance behavior. However, how descending neuromodulatory systems influence these distinct valence-associated circuits remained unknown.

Our findings reveal that OT achieves its dual prosocial functions through input-specific and output-specific circuit modulation. At the input level, OT selectively enhances inhibitory OTR neuron responses to noxious C-fiber activation (TRPV1+, MrgD+) while leaving responses to pleasant touch C-fibers (TH+, MrgB4+) relatively unchanged. At the output level, OT increases GPR83+ spinoparabrachial neuron responses to pleasant touch inputs while decreasing TACR1+ spinoparabrachial neuron responses to noxious inputs. This bidirectional modulation provides a circuit mechanism by which changes in internal state modulate spinal outputs, that in turn influence how supraspinal structures ultimately assign positive or negative valence to tactile experiences.

This represents a fundamental advance beyond earlier work showing that spinal OT modulates C-fiber and Aδ fiber inputs through GABAergic mechanisms. While those foundational studies established that the analgesic effect of spinal OT might act via enhanced inhibition in superficial dorsal horn, our work now reveals how this inhibition is precisely directed to suppress specific output pathways whose activation is associated with aversion while sparing or enhancing pathways whose activation is associated with reward. This provides the first molecular-circuit implementation of gate control theory with defined inputs, neuromodulators, and outputs that differentially influence behavioral outcomes associated with opposing valences.

### Volume transmission and laminar specificity

Consistent with previous anatomical studies, we observed no direct synaptic contacts between descending OT fibers and spinal OTR neurons, suggesting OT acts via en passant release and volume transmission. This mechanism is well-suited for the diffuse, state-dependent modulation we observe. However, our demonstration of a dorsoventral gradient in OTR neurotransmitter phenotype—from predominantly inhibitory in noxious-processing lamina I to predominantly excitatory in touch-processing lamina II inner—provides a structural basis for differential modulation. This anatomical organization, combined with the distinct C-fiber inputs to different laminae, explains how the same neuromodulator can simultaneously enhance inhibition of noxious processing while preserving excitatory signaling of pleasant touch.

Previous research using blind patch-clamp approaches in rat tissue provided important foundational insights. Breton and colleagues demonstrated that OT decreased IA potassium currents in lamina II interneurons, particularly those with bursting firing patterns, though these studies could not definitively identify neurotransmitter phenotypes. Our genetics-guided electrophysiological characterization reveals functional OTR expression in both excitatory and inhibitory spinal interneurons, with differential recruitment during distinct sensory contexts. This identification of mixed OTR populations, combined with our demonstration of their lamina-specific distribution and input-output connectivity to valence-encoding pathways, substantially advances understanding of how OT orchestrates spinal sensory processing.

### Evolutionary significance and species conservation

The joint evolutionary benefits of OT-mediated social behavior reinforcement and social touch-mediated pain relief likely contributed to conservation of this system across mammalian species. The ability of social touch to provide analgesia while simultaneously reinforcing social bonds would have conferred substantial survival advantages, particularly during injury, illness, or vulnerable states when social support is most critical. Our finding that mouse and human spinal OTR neurons show remarkably similar excitatory/inhibitory ratios despite 90 million years of divergence and dramatically different sensory ecologies suggests this balanced architecture represents a fundamental computational principle. Any deviation from this balance would bias the system toward either enhanced excitation or enhanced inhibition, destroying the flexibility required for context-dependent sensory reweighting. Disruptions in this precisely balanced excitatory/inhibitory architecture could provide a mechanistic framework for understanding altered touch processing in neurodevelopmental and psychiatric disorders—including autism spectrum disorder, anorexia nervosa, and depression—where touch aversion, hyperresponsivity to social touch, and reduced comfort from interpersonal contact represent core clinical features.

While both species show similar OTR organization, humans display broader OTR expression throughout the dorsal horn, which may reflect the greater importance of Aβ-mediated pleasant touch in humans. A recent study demonstrates that Aβ fibers contribute significantly to pleasant touch perception in humans93, and our human somatosensory evoked potential recordings—which activate Aβ fibers through electrical stimulation— showed spinal-level modulation following massage-induced OT release. This suggests the prosocial spinal OT mechanism extends beyond C-fiber modulation to encompass multiple touch modalities in humans.

### Clinical implications and therapeutic considerations

Our work has important implications for ongoing clinical trials of intrathecal oxytocin for chronic pain. While preclinical studies demonstrate robust analgesic effects of spinal OT activation, clinical trial results have been variable. Understanding OT’s multilayered roles throughout the body helps explain this complexity. OT acts peripherally19,94–97 though our characterization of OTR expression in adult mouse DRGs using the OTR-Venus and OTR-Tomato reporter mouse lines and RNAscope shows very low to undetectable levels (manuscript in preparation) and human DRG single cell RNA sequencing studies do not detect OTR expression in adult human DRGs98. OT also acts centrally in hypothalamic circuits controlling social homeostasis6, and in reward pathways including the VTA. Our findings reveal spinal OT modulation as another crucial layer in this coordinated system. This multilevel framework suggests a potential explanation for variable clinical outcomes: spinal OT alone may “prime” circuits by modulating their excitability and inhibitory tone, but tactile stimulation provides the sensory input necessary to fully engage these circuits. This is analogous to needing both a key and turning it—OT represents the key that changes circuit responsiveness across multiple anatomical sites, but appropriate sensory input is required to activate the primed pathways.

This framework suggests that optimal clinical outcomes might require combining intrathecal OT with structured tactile stimulation protocols (such as massage therapy) to maximally engage the reward-encoding pathways while suppressing pain-encoding pathways99,100. More broadly, our findings provide mechanistic legitimacy for massage as a treatment for chronic pain, moving beyond empirical observations to understanding the specific molecular and circuit mechanisms. Future work should identify which massage parameters most effectively trigger OT release and engage GPR83+ pathways while minimizing TACR1+ pathway activation.

### Broader implications for sensory processing

Our work demonstrates that the dorsal horn serves as an active computational hub where descending neuropeptidergic systems dynamically reconfigure sensory processing based on internal state. This aligns with emerging evidence that spinal circuits contribute to complex functions traditionally attributed to brain regions. The spinal integration of internal state and sensory information represents a previously unrecognized level of prosocial modulation in the nervous system.

The balanced representation of excitatory and inhibitory neurons among spinal OTR populations provides circuit flexibility necessary for bidirectional modulation. OT preferentially enhances inhibitory responses to achieve net analgesic effects while preserving touch reward circuits—a sophisticated integration occurring in circuits classically viewed as relay stations. Rather than passively transmitting sensory information, the dorsal horn actively integrates information about internal state to bias sensory experience toward prosocial outcomes through precisely organized circuits that can be modulated by descending neuropeptidergic systems in a context-dependent manner.

## Conclusion

We have uncovered evolutionarily conserved spinal neural circuits of touch reward and pain relief, revealing how changes in internal state following either rewarding touch or noxious experiences translate into circuit modifications that consistently promote prosocial outcomes. This OT-mediated feedback loop from affectively significant touch experiences to prosocial circuit priming represents a fundamental neuromodulatory mechanism that augments classical theories of sensory processing. The discovery that spinal OT differentially modulates valence-encoding spinoparabrachial pathways provides the first molecular-circuit implementation of gate control theory and establishes spinal OT circuits as a key mechanism linking social touch, pain relief, and adaptive behavior across mammalian species.

## Methods

### Animals

Mouse lines used in this paper include TH2a-CreER35 (Jax#025614), MrgB4-Cre (Jax#021077), MrgD-CreER101 (Jax#031286), TrpV1-Cre102 (Jax#017769), advillin-FlpO103, GAD2Cherry104 (Jax#023140), GAD67GFP105, Lbx1FlpO63, Cdx2-Cre106 (Jax#009350), GPR83GFP107, OTR-Cre108 (Jax#031303), OTR-Venus109,110, OTR-tdTomato (Jax#037580, RIKEN RBRC12289)111, OTR-flox109 (Jax#008471). For visualization and manipulation of target populations the following mouse lines were used: R26LSL-FSF-synaptophysin-GFP63,112 (RC::FPSit, Jax#030206), R26LSL-synaptophysin-tdTomato (Ai34, Jax#012570), R26LSL-tdTomato113 (Ai14, Jax#007914), R26LSL-FSF-ChR2-YFP108 (Ai80, Jax#025109). Transgenic mouse strains were used and maintained on a mixed genetic background (129/C57BL/6). All experiments were performed in adult mice unless otherwise stated. Experimental animals used were of both sexes. Housing, surgery, behavioral experiments, and euthanasia performed at Rutgers University were in compliance with Rutgers University Institutional Animal Care and Use Committee (IACUC; protocol #: 201702589). All mice used in experiments were housed in a regular light cycle room (lights on from 08:00 to 20:00) with food and water available ad libitum. All mice used in experiments were housed in a regular light cycle room (lights on from 07:00 to 19:00) with food and water available ad libitum.

### Tamoxifen treatment

Tamoxifen was dissolved in ethanol (20 mg/ml), mixed with an equal volume of sunflower seed oil (Sigma), vortexed for 5-10 min and centrifuged under vacuum for 20-30 min to remove the ethanol. The solution was kept at −80°C and delivered intraperitoneally for postnatal treatment.

### Immunohistochemistry and free-floating sections

Male and female P30-P37 mice were anesthetized with isoflurane and perfused transcardially (using an in-house gravity driven-perfusion system) with heparinized-saline (∼30 sec) followed by 15-20 minutes of 4% paraformaldehyde (PFA) in PBS at room temperature (RT). Vertebral columns were dissected and post-fixed in 4% PFA at 4°C for 2-24 hr. Sections were collected using a vibrating microtome (Leica VT1200S) and processed for immunohistochemistry (IHC) as described previously.210 Unless otherwise mentioned, transverse sections (50–60 μm thick) were taken from limb level lumbar spinal cord (L4-6). Free floating sections were rinsed in 50% ethanol (30 min) to increase antibody penetration, followed by three washes in a high salt PBS buffer (HS-PBS), each lasting 10 min. The tissue was then incubated in a cocktail of primary antibodies made in HS-PBS containing 0.3% Triton X-100 (HS-PBSt) for 48 at 4°C. Next, tissue was washed 3 times (10 minutes each) with HS-PBSt, then incubated in a secondary antibody solution made in HS-PBSt for 24 hr at 4°C. Immunostained tissue was mounted on positively charged glass slides (41351253, Worldwide Medical Products) and cover slipped (48393-195, VWR) with Fluoromount-G mounting medium (100241-874, VWR).

### Fluorescence in situ hybridization (FISH; RNAscope)

Mice were anesthetized with isoflurane and perfused transcardially as described above. Spinal cord and dorsal root ganglia were dissected and post-fixed in 4% PFA at 4°C for 2-24 hr. Samples were cryosectioned at 20 μm and processed using the RNAscope Multiplex Fluorescent v2 Assay (Advanced Cell Diagnostics, 323110). Tissue was placed in 1xPB for 5 mins at room temperature then dehydrated through successive EtOH steps (50%, 70%, 100%, 100% – 5 min) and then dried at room temperature. Tissue was then treated with hydrogen peroxide for 10 min at RT, washed in water, treated with target retrieval reagents (322000) in a steamer for 5 min, and then treated with 100% EtOH for 3 min at RT. Tissue was then treated with protease III (322337) for 8 min at 40°C before being rinsed with water. Probes were hybridized for 2 hrs at 40°C in a humidified oven, rinsed in wash buffer, and then stored overnight in 5x saline sodium citrate. After rinsing with wash buffer, a series of incubations was performed to amplify and develop hybridized probe signals. Briefly, AMP1 (323101), AMP2 (323102), and AMP3 (323103) were successively applied for 30 min, 30 min, and 15 min at 40°C in a humidified oven, respectively, with a buffer wash in-between. For each channel, HRPC1 (323104) and/or HRPC2 (323105) and/or HRPC3 (323106) were applied for 15 min at 40°C in a humidified oven followed by a buffer wash, then treated using TSA Plus Fluorophores (PerkinElmer; NEL741001KT, NEL744001KT, and NEL745001KT), followed by 15 min of HRP blocker (323107) at 40°C in a humidified oven. Slides were then mounted with Fluoromount-G mounting medium (100241-874, VWR) with DAPI.

### Image acquisition and analysis

Images were captured with a Zeiss LSM 800 confocal. Images for cell counts were taken with 10x or 20x air objectives, and images of synaptic contacts were taken using a 40x oil objective. ImageJ (cell count plug-in) was used for colocalization analysis of cell bodies. Imaris (spot detection plug-in) was used for synaptic analysis, quantification of synaptic terminals, and generation of contour density plots. Only puncta apposing a postsynaptic marker Homer1 for excitatory synapses were included for analysis. To generate contour density plots, Imaris spot detection plug-in was used to detect the spatial coordinates of cell bodies/synaptic terminals. Following detection, coordinate values were corrected and expressed relative to common reference points (central canal, dorsal, lateral and ventral edges of the gray matter). Corrected coordinates were used to obtain two-dimensional kernel density estimation utilizing the MATLAB ‘ksdensity’ function. Kernel density estimations were graphically displayed as contour plots, using the MATLAB ‘contourf’ function.

### c-Fos induction

Mice were transferred to a holding area adjacent to the testing room at least 7 days prior to the experiment, to decrease stress during the task. 8-12 weeks old mice were either anesthetized with isoflurane, their hairy back skin was gently brushed with a soft blush brush similarly to the brush-CPP assay (twice for 100 sec 5 min apart), kept under anesthesia for an hour then perfused for immunohistochemistry, or stimulated with noxious heat until a nocifensive reflex was observed then immediately anesthetized with isoflurane for an hour and perfused for immunohistochemistry, or had spared nerve injury surgery performed then three weeks later were anesthetized with isoflurane and perfused for immunohistochemistry. Transverse sections (50-60 μm thick) were taken from lumbar spinal cord and stained. A Zeiss LSM 800 confocal was used to acquire 20x images. Images were then used for quantification of c-Fos expressing neurons.

### Spinal cord injections

Mice were anesthetized with isoflurane (5% induction, 1.5-2% maintenance). A small incision was made in the skin and underlying musculature and adipose tissue was teased apart to reveal the vertebral column. Vertebral surfaces were cleaned with fine forceps to reveal the dorsal surface of the spinal cord. Viruses were injected into 3-4 sites in the L4-L6 spinal cord through a pulled glass needle, ∼150 nL per site. Following injections, the overlying muscle and the skin incision was sutured. Mice were allowed to recover from anesthesia and returned to their home cages. 3-4 weeks following the injection, mice were perfused, and their tissue was processed for immunohistochemistry.

### Spared nerve injury (SNI)

Mice were anesthetized and the hair surrounding the hindlimb was shaved. An incision was made on the right left below the knee, and another incision was made in the muscle to reveal the sciatic nerve. A ligature using silk suture (Braintree Scientific, SUT-S 103) was made around the peroneal and tibial nerve branches, leaving the sural nerve intact. A 2 cm portion of the peroneal/tibial nerve was cut and removed. Special care was taken to ensure that the sural nerve was not damaged during the procedure. Surrounding musculature was sutured around the nerve, and then the wound was secured using surgical wound clips (Kent Scientific, INS750344-2). Tactile hypersensitivity was determined using von Frey testing, and animals exhibiting any paralysis or total desensitization were excluded from experiments.

### Behavior

Male and female mice 12-26 weeks of age were used for behavioral tests.

### Mechanical sensitivity

Paw withdrawal thresholds in response to mechanical stimulation were measured with two calibrated von Frey filaments (0.07 and/or 0.4 g, Stoelting Co., Wood Dale, IL) as described (Jia et al, 2022; Mao et al, 2019; Wei et al, 2021; Wen et al, 2023; Yang et al, 2021)114–118. Briefly, mice were placed in a Plexiglas chamber on an elevated mesh screen. Each von Frey filament was applied to the plantar sides of both hind paws 10 times. A quick withdrawal of the paw was regarded as a positive response. The number of positive responses among ten applications was recorded as percentage withdrawal frequency (number of paw withdrawals/10 trials) × 100 = % response frequency.

### Heat sensitivity (plantar test/Hargreaves)

Paw withdrawal latencies in response to noxious heat stimulation were examined as published (Jia et al, 2022; Mao et al, 2019; Wei et al, 2021; Wen et al, 2023; Yang et al, 2021)114–118. Briefly, mice were placed in a Plexiglas chamber on a glass plate. A beam of light emitted from a hole in the light box of a Model 336 Analgesia Meter (IITC Inc. Life Science Instruments, Woodland Hills, CA) was applied to the middle of the plantar surface of each hind paw. A quick lift of the hind paw was regarded as a signal to turn off the light. The length of time between the start and the stop of the light beam was defined as the paw withdrawal latency. For each side, three trials at 5-min intervals were carried out. A cut-off time of 20 s was used to avoid tissue damage to the hind paw.

### Conditioned Place Preference/Aversion (CPP/CPA)

The CPP/CPA test was carried out as described with minor modifications (Jia et al, 2022; Mao et al, 2019; Wei et al, 2021; Wen et al, 2023; Yang et al, 2021)114–118. Briefly, an apparatus with two Plexiglas chambers connected with an internal door (Med Associates Inc., St. Albans, VT) was used. One of the chambers was made of a rough floor and walls with black and white horizontal stripes, and another one was composed of a smooth floor and walls with black and white vertical stripes. Movement of the mice and time spent in each chamber were monitored by photobeam detectors installed along the chamber walls and automatically recorded in MED-PC IV CPP software. Mice were first preconditioned for 30 min with full access to two chambers to habituate them to the environment. At the end of the preconditioning phase, basal duration spent in each chamber was recorded within 15 min to check whether animals had a preexisting chamber bias. Mice spending more than 80% or less than 20% of total time in any chamber were excluded from further testing. The conditioning protocol was performed for the following 3 days with the internal door closed. The mice first received an intraperitoneal injection of clozapine N oxide (CNO) coupled with either an intrathecal saline injection (5 µl) paired with one conditioning chamber in the morning, or, six hours later, an intrathecal lidocaine injection (0.8% in 5 µl of saline) paired with the opposite conditioning chamber in the afternoon. Lidocaine at this dosage did not affect motor function. The injection order of saline and lidocaine was switched every day. On the test day, at least 20 h after the conditioning, the mice were placed in one chamber with free access to both chambers. The duration of time that each mouse spent in each chamber was recorded for 15 min. Score differences were calculated as test time-preconditioning time spent in the lidocaine chamber.

### Homecage single-housing and pre-conditioning stroking

Before stroking, mice were single-housed in the homecage for one week, with non-soft enrichment: a wooden block and a glass or hard plastic hut. Mice were pre-conditioned with gentle stroking in the homecage. The lid and grid were removed 30 min before stroking for acclimation. Gentle stroking was performed by a trained experimenter who was blind to mouse genotype/strain information/virus injected. The hairy back skin of mice was gently stroked with a hand-held soft brush (e.l.f. Cosmetics Blush Brush), moving from the nape of the neck to the lumbar enlargement region (T11∼L2) at a constant speed and force as previously described (21). The stroking was repeated daily for four trials (one trial includes 100-s stroking twice and 5-min rest in between). The brush was not repeatedly used for different animals to avoid odor cross-interaction.

### Gentle brush conditioned place preference

To assess the valence of gentle stroking, we employed an unbiased two-chamber CPP apparatus (two compartment CPP package Med Associates, MED-CPP2-3013-2) consisting of two equally sized chambers separated by a small connecting opening. The two chambers are differentiated by distinct visual and tactile cues, including black or white walls and a grid rod or mesh floor. The CPP protocol consists of three phases: pre-conditioning phase to measure the baseline, conditioning phase and the test phase.

For the pre-conditioning phase, mice were placed in a random chamber and allowed to freely explore the CPP box for 15 min with the door open. The time spent in each chamber was measured by infrared beams at the bottom of the chambers and calculated by Med Associates software. This was done over three days to ensure a clear assessment of the baseline preference.

For the conditioning phase, 26 conditioning sessions were performed, two sessions per day over 13 days. One session includes one chamber paired with stroking and the other without, on consecutive days, interchangeably in the morning or the afternoon. The conditioning time includes 30 min acclimation, followed by 15 min stroking that consists of 2 stroking trials, each with 100-s stroking and 5-min rest. The stroking is applied in the chamber where mice spent the least amount of time at baseline. For the non-stimulus conditioning, mice were subjected to the same procedure as described above except for stroking trials.

For the test phase, mice were allowed to freely explore the CPP box for 15 min on the test day. The time mice spent in each chamber was recorded and analyzed by Med Associates software.

### Tactile Pre Pulse Inhibition (tactile PPI)

The response of mice to tactile startle stimuli was measured using a San Diego Instruments startle reflex system (SR-LAB Startle Response System) (Orefice et al., 2016). In brief, for tactile PPI air puffs were administered to the back of the mouse to assess hairy skin sensitivity. A 1.5 PSI air puff prepulse stimulus strength was chosen because control animals of this particular Bl6/FVB mix showed a minimal, but statistically significant response to the stimulus alone, compared to baseline movement in the chamber without any stimulus (average response in controls, 8.19 ± 1.39%). Each mouse was placed in the chamber for a 5 min acclimation period, during which constant background noise of broadband white noise was presented. Background noise for the tactile PPI testing sessions was 75 dB, to ensure that the animal could not hear the air puff prepulse. The prepulse intensity remained constant (1.5PSI, 50ms), and the ISI varied from 50 ms to 1 s in duration. Whole body flinch, or startle reflex, was quantitated using an accelerometer sensor measuring the amplitude of movement of the animal, within the cylindrical holder.

### Mouse electrophysiology experiments — Stereotaxic injection

Mice aged 6–8 weeks were deeply anesthetized with isoflurane and then head-fixed in a stereotaxic frame (David Kopf Instruments). A small craniotomy was made over the PBNL injection site, using these coordinates relative to the bregma (mm): AP: −5.2 – −5.0, ML: ± 1.4 – 1.6; DV: −2.8 – −3.0. Borosilicate pipettes with 5–10-µm-diameter tips were backfilled with virus (pENN.AAV.CB7.CI.mCherry.WPRE.RBG, # 105544, addgene), and a volume of 150–250 nl was pressure-injected using a Nanoject III (Drummond) every 30 s. The pipette was left in place for an additional 5 min, allowing time to diffuse away from the pipette tip, before being slowly retracted from the brain. For both retrograde and viral labeling, animals were housed for 2–3 weeks before slicing. Acute transverse spinal cord slices were used for whole-cell patch-clamp recordings of retrogradely labeled Gpr83+ and putative Tacr1+ SPB neurons.

### Slice preparation

Mice aged 8–12 weeks were anesthetized with an intraperitoneal injection of a lethal dose of ketamine and xylazine and perfused intracardially with an ice-cold external solution containing the following (in mM): 250 sucrose, 25 NaHCO3, 1 NaH2PO4, 10 glucose, 2.5 KCl, 6 MgCl2, 1 CaCl2 (295–305 mOsm) and bubbled with 95% O2/5% CO2. Coronal slices (300 mm thick) were cut on a Compresstome VF-200 slicer (Precisionary Instruments Inc., Greenville, NC, USA), then immediately incubated in oxygenated (95% O2/5% CO2) standard artificial cerebrospinal fluid (ACSF) containing the following (in mM): 120 NaCl, 25 NaHCO3, 1.4 NaH2PO4, 21 glucose, 2.5 KCl, 2 CaCl2, 1 MgCl2 for at least hour at room temperature (24°C-25°C) until transferred to the recording chamber.

### Electrophysiology

All recordings were made at room temperature (22–24°C) and neurons were identified by using a LEICA DM6000 FS microscope equipped with infrared differential interference contrast, a CCD camera, and fluorescence. For all electrophysiology recordings, slice was transferred to a recording chamber and continually superfused with ACSF bubbled with 95% O2/5% CO2 to achieve a pH of 7.3–7.4. All recordings were performed using a Multi-Clamp 700B amplifier (Molecular Devices, Sunnyvale, CA, USA) and a Digidata 1440A digital interface (Axon Instruments, Union City, CA, USA). For whole-cell recordings, a K+-based pipette solution containing 142 mM K+-gluconate, 10 mM HEPES, 1 mM EGTA, 2.5 mM MgCl2, 4 mM ATP-Mg, 0.3 mM GTP-Na, 10 mM Na2-phosphocreatine (295 mosmol, pH 7.35) or a Cs+-based pipette solution containing 121 mM Cs+-methanesulfonate, 1.5 mM MgCl2, 10 mM HEPES, 10 mM EGTA, 4 mM Mg-ATP, 0.3 mM Na-GTP, and 10 mM Na2-Phosphocreatine (295 mosmol, pH 7.35) was used. Membrane potentials were not corrected for liquid junction potential (experimentally measured as 12.5 mV for the K+-based pipette solution and 9.5 mV for the Cs+-based pipette solution).

For voltage clamp experiments, cells were held at −60 mV and excitatory postsynaptic currents (EPSCs) are recorded. Resting membrane potential (RMP) was recorded in current-clamp mode with no holding current (I = 0), and action potential (AP) discharge was recorded in current-clamp evoked by depolarizing current steps (0– 140 pA, 1000 ms). For pharmacology experiments, the baseline synaptic currents were recorded for at least 5 min in the absence of any drug. The drugs were then added to the ACSF at the following concentrations: Oxytocin (2 mM), TTX (1 μM), DNQX (10 μM), and 4-AP (1.5 mM). The blue light was emitted from a collimated light-emitting diode (LED) of 470 nm. The LEDs were driven by a LED driver (Mightex) under the control of an Axon Digidata 1440A Data Acquisition System and Clampex 10.5. Light was delivered through the reflected light fluorescence illuminator port and the 40× objective.

For light pulse stimulation, pulse duration (0.1–10 ms) and intensity (2.3–24mW/mm2) were adjusted for each recording to evoke reliable synaptic EPSCs. Light pulses were delivered at 30 s interstimulus intervals. The blue light source (Polygon 400, Mightex, Pleasanton, CA, USA) used to stimulate channel rhodopsin at 470 nm was mounted on the side of the microscope to a dichroic mirror located between the binoculars and the objective, allowing the light to be directed toward the slice through the 40 × objective. The spatial illumination pattern on the slice was controlled by Dynamic Spatial Illuminator software (Mightex) in conjunction with a digital camera. The duration and intensity of the light stimuli were set in BioLED source control module software (Mightex). The light stimulus was controlled in pClamp 10.5 software (Axon Instruments/Molecular Devices, Sunnyvale, CA, USA) using a digital output channel to connect the Digidata 1440 A digitizer to the BioLED controller.

### Data analysis

Cell capacitance, current amplitude, latency, and jitter were analyzed using Clampfit 10.5. Intrinsic properties were determined as follows. Input resistance was calculated from the steady-state voltage during a −20 pA, 1000 ms current step. The maximum firing rate was determined by increasing the current injection in a stepwise fashion until the firing rate plateaued. Rheobase was determined by the average minimum current injection that elicited firing. For experiments with a single optogenetic stimulation, the PSC amplitude was measured as the average value across 1 ms around the peak subtracted by the average 100 ms baseline value prior to the stimulation.

### Mouse fiber photometry of spinal cord-projecting OT neurons — Intraspinal viral injections and optic fiber implantation

7-10 week old females were pretreated with 5mg/kg oral or intraperitoneal meloxicam before surgery. They were anesthetized in a chamber with 3% isoflurane and kept anesthetized with 1.5-2% isoflurane. Laminectomies were performed to expose either cervical, thoracic, or lumbar spinal cords. Bilateral injections of 150nl into two to three spinal segments for a total 900 nl of retrograde Cre-dependent calcium indicator expressing AAV viruses were directly injected in the spinal cord of OT-Cre mice using pulled glass pipettes (Wiretrol II, Drummond) and a microsyringe pump injector (UMP3, World Precision Instruments).

One week later, we implanted the optic fiber (Doric, MFC_200/230-0.57_6mm_MF1.25_FLT) 200μm above the paraventricular nucleus of the hypothalamus. Three skull screws were inserted in the skull to stabilize the implant. A small amount of Metabond cement (Parkell) was used to bond the fiber, skull, and skull screws. Dental cement was applied on top of dried Metabond to create a stable implant structure. Mice were given 1 week to recover from implant surgery and 2 weeks to recover from injection and implant combined surgeries before behavioral testing. Meloxicam was administered 24 hours after surgery during post-operative monitoring.

### Fiber photometry

Zirconia sleeves (Doric) were used to connect the optic fiber implant to the patch cord. Signals were recorded using a real-time processor (RZ10X, TDT) and extracted in real time using Synapse software (TDT). A 465nm LED was used to excite the virally encoded calcium sensor while a 405nm LED was used as isosbestic to measure and control for changes in fluorescence due to photobleaching and movement artifacts.

For fiber photometry during gentle brushing, mice were habituated to gentle brushing for 3 days prior in their home cage. During the recording session, mice were left in their home cage and, after 5 min baseline recording, gentle brushing of back hairy skin was applied similarly to the gentle brush CPP assay: twice for 100 sec with a 5 min pause in between stimulations. For fiber photometry during the dynamic hot plate assay, mice were placed on the hot plate surrounded by a plexiglas cylinder to allow the cord to connect from the head to the computer. After 5 minutes baseline recording, the heat ramp was started, with temperature increasing from room temperature to 52 degrees Celsius followed by a 10 min cool down period.

### Mouse 2P ex vivo calcium imaging experiments — Animals

All animal procedures followed the ethical guidelines of the International Association for the Study of Pain and ethical guidelines of the Directive 2010/63/UE of the European Parliament and of the local ethical committee (authorization project 11837-2017101816028463V5), on the protection of animals used for scientific purpose. Double heterozygote VGLUT2 cre/+ ; R26 lsl-GCamp6s/+ and VGAT cre/+ ; R26 lsl-GCamp6s/+ mice were used for Ca2+ imaging. Both male and female mice (4- to 8-weeks old) were used for this study. No significant sex differences were observed and data from each sex were pooled. Mice were kept on an inverted 12:12 light/dark cycle (light OFF 7am-7pm) in micro-isolator caging racks (Tecniplast) with food and water provided ad libitum.

### Spinal cord ex vivo somatosensory preparation

Mice were deeply anesthetized with an i.p. injection of 100 mg/kg ketamine and 20 mg/kg xylazine. The back of the mouse was shaved and a dorsal laminectomy was performed. The spinal cord with dorsal root ganglions still attached was quickly harvested and transferred to a dissection dish perfused with an ice-cold (4°C) solution containing (in mM) 194 sucrose, 2.0 KCl, 7.0 MgCl2, 26 NaHCO3, 1.15 NaH2PO4, 11 D-glucose, and 0.5 CaCl2 (290-300 mOsm.kg-1), pH 7.4 bubbled with 95% O2 and 5% CO2. The dura matter was removed, and the spinal cord with the dorsal roots attached was embedded into 3% low-melting agarose (Fisher Scientific 10377033). 550 µm transverses slices of spinal cords with the L4 or L5 DRG still attached were obtained using a vibroslice (VT1200 S; Leica Microsystèmes SAS, Nanterre, France). Slices were left to recover for at least 1 hour in an artificial cerebrospinal fluid (aCSF) containing (in mM): 128 NaCl, 3 KCl, 2.5 CaCl2, 1.5 MgSO4, 0.6 NaH2PO4, 25 NaHCO3, 10 glucose (300-310 mOsm.kg-1), pH 7.4, bubbled with 95% O2 and 5% CO2. After recovery, slices were transferred to a recording chamber (volume ≈3 ml) and continuously perfused at 3 ml/min with oxygenated aCSF at room temperature (≈23°C).

### Drugs application of spinal cord slices

Oxytocin (Sigma-Aldrich #04375) was incrementally bath applied at a final concentration of 0.01, 0.1 and 1 µM, respectively. To determine the total number of excitatory or inhibitory dorsal horn potential responding neurons, GABAergic and glycinergic spinal cord disinhibition was performed at the end of each recording as previously described (Torsney and MacDermott, 2006) using bath application of 10 μM bicuculline (Sigma-Aldrich #14343) and 1 µM strychnine (Sigma-Aldrich #S8753). In each case, recordings began 30 min after the onset of drugs application, and during continuous perfusion of drugs at the appropriate concentration.

### Dorsal root electric stimulations

Dorsal roots were stimulated using a suction electrode and a A365 stimulus isolator (World Precision Instrument) as previously described (Peirs et al., 2015)119. Roots were stimulated at 2 Hz with a pulse duration of 0.1 ms. Afferent fibers were stimulated at different intensities to recruit Aβ, Aδ and C fiber with 25, 100 and 500 μA pulses, respectively.

### Two-photon calcium imaging

Ca2+ imaging was performed using an upright Zeiss Axio Examiner Z1 microscope and the Zen black acquisition software (Carl Zeiss). Images were acquired using a Plan-Apochromat 20x/1.0, water-immersion objectives and a Multiphoton LSM 7 MP (Carl Zeiss). For each slice, four individual focal planes of 425×425 µm window, separated by 15-20 µm intervals were illuminated with a sapphire Mai Tai laser (Spectra-Physics, USA) at 920 nm. The border of the scanning area was set near the dorsal border between the white mater and the grey matter of the DH to allow the visualization of DH neurons located in both superficial and deeper laminae. For electric stimulations, baseline fluorescence intensity was first acquired during 15 seconds, then the dorsal roots were stimulated for 10 seconds, and fluorescence was recorded for an additional 15 seconds.

### Calcium imaging analysis

At the end of each experiment, the maximum number of responding neurons was assessed following suprathreshold stimulations of the dorsal root with 500µA, under spinal disinhibition. Only neurons that responded to such stimulation were kept further for analysis. Images were exported into MATLAB (MathWorks) for analysis with a custom-made toolbox. Using this toolbox, images from each recording were x-y motion corrected, and recordings where a significant displacement in the Z leading to the loss of the focal point were excluded. In turn, the ΔF/F and the standard deviation of fluorescence over time for each focal plane were analyzed individually, and ROI manually drawn around clearly identified cell bodies. ΔF/F was calculated as ΔF/F =( F(t) - F0)/F0 where F(t) is the fluorescent value at a given time and F0 as the mean fluorescence during baseline recording. Neurons were considered responders when the ΔF/F during the stimulation window exceeded 3 times the standard deviation of the mean fluorescence during baseline and with at least a peak amplitude 20% higher than any other peak during the baseline period. Data was calculated as percentage of total DH neurons responding to dorsal root stimulations at 500µA under spinal disinhibition. For segmentation of neurons receiving Aβ, Aδ or C-fibers inputs, any cell that stopped responding during the ramp of electric stimulations were excluded from this analysis to avoid potential false negative interpretation of sensory inputs.

### Statistical analysis

Statistical analyses of Ca2+ imaging data were performed using GraphPad Prism 9.0. Contingency analysis was performed using Fisher’s exact tests. Differences were considered significant if p<0.05. Figures were made using CorelDRAW X7.

### Human psychophysical studies — Participants

A total of thirty-seven female participants living in Sweden were included in the study, however, five participants only took part in one session and dropped out of the study, the final amount attending two laboratory visits included were 32 females (mean age= 24,72, SD= 4,08), for an approximate duration of 2 hours. Two laboratory visits were performed, an experimental visit and a control visit. The order of the visits was counterbalanced, we included 19 participants that started with the experimental visit and 13 with the control visit. Most participants had a small or absent menstrual cycle due to being on hormonal medication, 20 subjects were on hormonal medication (11 taking contraceptive pills majority with no placebo, 9 were on hormonal IUD). Efforts were made to have participants come in for their second visit at approximately at the same date of cycle than done previously, only 2 participants who were not on any hormonal medication could only re-join the study at a different cycle stage with a difference of 15 and 21 days of difference from their previous evaluation.

Participants were recruited through social media advertisements and were screened before inclusion. This sample size was based on a power calculation for a two-phase cross-over comparison with 80% power to detect an effect size of Cohen’s D ≥ 0.6 at alpha = 0.05, i.e., a medium—large effect size. Inclusion criteria included: pre-menopausal, non-pregnant female, between 18-40, willing to receive a massage and finding it pleasant. Exclusion criteria included: any clinically significant medical condition, any current clinically significant psychiatric or motor-sensory problems, unwilling to receive or find it disconformable to receive unpleasant stimulation to the arms. Participants received 1000 Swedish kronor as reimbursement for both visits (approx. 50 taxable euros for each visit). All participants provided written consent before study participation. The project was approved by Swedish Ethic authority (dnr 2023-04239-01).

### Study Procedure

After inclusion, participants attended two lab visits: one experimental (massage) and one control (questionnaires), in a counterbalanced order. At the start of each visit, participants rinsed their mouths with water and washed their hands to minimize contamination of saliva samples. They then sat in a massage chair, regardless of the visit type, and the electrophysiology setup commenced. Electrodes were attached to the scalp, spine, Erb’s point, and chest (ECG).

Each visit began with the collection of the first saliva sample, followed by a resting-state EEG1/ECG1 recording (EEG1). Next, participants’ personal thresholds for electrical and thermal stimulation were determined, which were kept consistent across both sessions. Somatosensory evoked potentials (SEP) were measured by electrically stimulating the ulnar nerve, and cortical contact heat evoked potentials (CHEP) were assessed by stimulating the forearm using a temperature probe (QST.Lab), and are reported elsewhere. After each SEP and CHEP stimulation, participants provided subjective ratings of unpleasantness and pain.

The intervention phase lasted approximately 30 minutes and involved either a soft, hand-delivered massage designed to promote relaxation (experimental visit) or the completion of questionnaires on an iPad (control visit). Post-intervention, the second saliva sample was collected, followed by subjective ratings of relaxation. Resting-state EEG2/ECG2 was recorded next—during the massage in the experimental visit, and with participants sitting with eyes closed in the control visit. SEP2 and CHEP2 were performed using the same stimulation thresholds as earlier, followed by a third SEP (SEP3) assessment. After removing all electrophysiology equipment, the third and final saliva sample was collected.

In summary, the outcome measures for each visit included: 1) Oxytocin (3 samples) and 2) cortisol (3 samples) concentrations in saliva, 3) SEP (3 times) with subjective ratings, 4) CHEP (2 times) with subjective ratings, 5) ECG (2 times), 6) PANAS questionnaire (1 time).

### Data processing — Salivary Oxytocin Collection and Assay

Saliva samples were collected via passive drool at three timepoints: baseline, post-intervention, and at the end of each visit. A total of 207 saliva samples were collected, with no samples classified as quantity not sufficient (QNS). Samples were stored at −80°C prior to shipment on dry ice to the Salimetrics SalivaLab (Carlsbad, CA) for analysis. Upon arrival, the samples were thawed to room temperature, vortexed, and centrifuged for 20 minutes at approximately 3,500 RPM (1,500 x g) immediately before testing. Oxytocin levels in the saliva were assayed in triplicate using a proprietary electrochemiluminescence method developed and validated by Salimetrics specifically for salivary oxytocin detection. Each assay used 25 μl of saliva per determination. The assay’s sensitivity threshold was 8.0 pg/mL, with a standard curve range of 6.4–1600 pg/mL. The average intra-assay coefficient of variation was 20.48%, while the average inter-assay coefficient of variation was 3.90%, ensuring accuracy and repeatability according to Salimetrics’ criteria and NIH guidelines for reproducibility in salivary bioscience research.

### Salivary Cortisol Collection and Assay

Saliva samples for cortisol analysis were collected using the SalivaBio Oral Swab (SOS) (Item No. 5001.02). The SOS was placed under the tongue for 3 minutes at baseline, post-intervention, and at the end of each visit (experimental and control). The SOS offers ease of use and helps filter large macromolecules and other particulates, improving assay results. Due to the consistency across lots, the SOS was ideal for this longitudinal, multi-participant study. After collection, the swabs were stored in a refrigerator until they were centrifuged to remove the swab and extract the saliva. Following centrifugation, the saliva samples were initially stored at −20°C, as the collection tubes used were not cryotubes. The samples were then transferred to −80°C for long-term storage, but for no longer than 3 months.

Cortisol levels were analyzed using a competitive enzyme-linked immunosorbent assay (ELISA). In this assay, cortisol from the samples competes with cortisol conjugated to horseradish peroxidase for antibody binding sites on a microtitre plate. After incubation, unbound components were washed away, and bound cortisol-enzyme conjugate was measured through a reaction with tetramethylbenzidine (TMB), producing a blue color. The reaction was stopped with an acidic solution, turning the solution yellow. Optical density was measured at 450 nm using a plate reader, with the amount of cortisol-enzyme conjugate inversely proportional to the cortisol concentration in the samples. Salivary cortisol data further characterize the endocrine context and will also be reported in another manuscript.

### Hormonal Analysis

Hormone outcomes were analyzed using repeated-measures models with Treatment (experimental vs control) and Time (three repeated measurements) as within-subject factors. Analyses were performed on participants with complete data across the Treatment × Time design (complete-case analysis). Prior to inferential testing, outliers exceeding ±2 SD from the condition distribution were excluded. For repeated-measures ANOVA, the sphericity assumption was evaluated with Mauchly’s test; when violated, Greenhouse–Geisser corrected degrees of freedom were reported. Tests were two-sided with α = 0.05. Effect sizes are reported as partial eta squared (ηp²). When significant main effects were observed, post hoc pairwise comparisons were conducted using Tukey’s test, with family-wise error (FWE) correction applied across the full set of comparisons.

Oxytocin (Fig 1.c): Oxytocin concentrations changed significantly over time (repeated-measures ANOVA; main effect of Time, Greenhouse–Geisser corrected: F(1.758, 38.686) = 9.307, p < 0.001, ηp² = 0.297). There was no main effect of Treatment (F(1, 22) = 0.132, p = 0.719, ηp² = 0.006). The Treatment × Time interaction did not reach the prespecified significance threshold, but showed a trend (Greenhouse–Geisser corrected: F(1.579, 34.729) = 2.820, p = 0.085, ηp² = 0.114; uncorrected p = 0.070). Analyses were conducted in participants with complete data across the Treatment × Time design (n = 23), after excluding outliers (> ±2 SD). During massage session, salivary oxytocin increased from pre-massage to both immediate and late post-massage time points (mean pre = 9.76 pg/mL; mean post1 = 15.84 pg/mL; mean post2 = 15.069 pg/mL; n = 23), as confirmed by Tukey-FWE corrected post hoc comparisons following the repeated-measures ANOVA (p = 0.007 and p = 0.026, respectively).

Salivary cortisol levels were analyzed using a 2 (Treatment: massage, control) × 3 (Time: baseline, post, post2) repeated-measures ANOVA after removal of outliers exceeding ±2 SD. A significant main effect of Time was observed (Greenhouse–Geisser corrected: F(1.333, 33.333) = 11.554, p < 0.001, ηp² = 0.315), indicating changes in cortisol levels across the session. There was no main effect of Treatment (F(1, 25) = 0.446, p = 0.510, ηp² = 0.016) and no Treatment × Time interaction (Greenhouse–Geisser corrected: F(1.242, 31.038) = 1.258, p = 0.281, ηp² = 0.048). Post hoc Tukey-corrected pairwise comparisons revealed significant reductions in cortisol from baseline to post-session time points within the massage condition (baseline vs post1: p = 0.047; baseline vs late post2: p < 0.001). No significant differences were observed between post and late post time points, nor between treatment conditions at matched time points (all p > 0.05).

### Positive and Negative Affect Schedule (PANAS)

The Positive and Negative Affect Schedule (PANAS) questionnaire (Watson et al., Soc Psychol 1988) was administered once during each visit. Participants completed the PANAS to assess their affective state, measuring negative affect, at a single time point during the experimental (massage) and control visits. Because positive affect items largely index arousal (e.g. Interested, Excited, Strong, Enthusiastic, Alert), primary analyses focused on negative affect (Distressed, Upset, Guilty, Scared, Hostile, Irritable, Ashamed, Nervous, Jittery, Afraid). Scores were derived by summing the corresponding PANAS items. Affect ratings obtained during the massage and control visits were compared using paired-samples t-tests (two-sided, α = 0.05).

Negative affect scores were significantly lower during the massage visit compared to the control visit (paired t-test, t(29) = 4.339, p < 0.001), indicating reduced negative affect during massage (control: mean = 20.23, SD = 5.90; massage: mean = 14.65, SD = 4.06).

### Somatosensory evoked potentials (SEP)

A stimulation electrode was placed on the inside of the left forearm, targeting the ulnaris nerve. Subjects were asked to close their eyes and relax, stay still but not fall asleep during stimulation. Recording electrodes were placed on the Erb’s point (targeting the brachial plexus), the C4 cervical level, and on C4, CZ, and FZ scalp positions. According to previously reported clinical neurophysiology protocol (Boehme PNAS, 2019), a maximum of 300 nonpainful pulses at a maximum of 100 mA. Stimulation intensity was individually adjusted to the minimum current for each participant necessary to evoke a ring and little finger twitch at 1 Hz resulting in a length of less than 5 min for each time-point (SEP1, SEP2 and SEP3). Electrode skin impedance was always less than 8 kΩ. Data were acquired for 100 ms after each pulse using a NicoletEDX system with an AT2+6 amplifier (Carefusion) and recorded and analyzed using Synergy 20.0 (Carefusion). Recordings were online referenced to Fz and bandpass filtered (2 Hz to 2 kHz), the amplifier range was 5 mV, and the display sensitivity was 20 μV per division. Waveforms were averaged over the total amount pulses for each recording electrode and over the two runs per condition and analyzed with regard to amplitude and latency (N13 cervically, N20 cortically). Baseline to peak amplitude was calculated automatically, with the baseline defined as the value right before the average waveform and with automatically selected peaks (Desmedt JE, Cheron G, Clin Neurophysiology, 1980), which were inspected individually and manually adjusted. Values were compared using repeated measures ANOVA and using Tuckey as postdoc two pair contrasts and FWE to correct for multiple comparisons.

Subjective discomfort ratings were collected. Following each SEP recording (SEP1, SEP2, SEP3), participants rated perceived discomfort in response to the electrical stimulation on a numeric scale ranging from 1 (not uncomfortable) to 10 (extremely uncomfortable). Discomfort ratings were analyzed using a repeated-measures ANOVA with Treatment (massage, control) and Time (SEP1, SEP2, SEP3) as within-subject factors. When appropriate, Tukey post hoc comparisons with family-wise error (FWE) correction were performed to explore pairwise differences.

(fig 3). Discomfort ratings. Subjective discomfort ratings changed across SEP recordings, as indicated by a significant main effect of Time (F(2, 58) = 7.472, p = 0.001, η² = 0.038). There was no main effect of Treatment (p = 0.747) and no Treatment × Time interaction (p = 0.131). Post hoc Tukey FWE-corrected comparisons revealed a significant change in discomfort ratings between the second and third SEP recordings within the massage condition (p = 0.010), whereas other pairwise comparisons were not significant.

(Fig 7) Spinal SEP latency. Cervical SEP (N13) latency changed significantly across time (F(2, 48) = 8.831, p < 0.001, η² = 0.107), with no main effect of Treatment and no Treatment × Time interaction. Post hoc Tukey comparisons indicated a significant increase in SEP latency from baseline (t1) to post-massage (t2) within the massage condition (t1: mean = 13.76 ms; t2: mean = 14.07 ms; p = 0.038), whereas no other comparisons reached significance.

**Figure 7:**
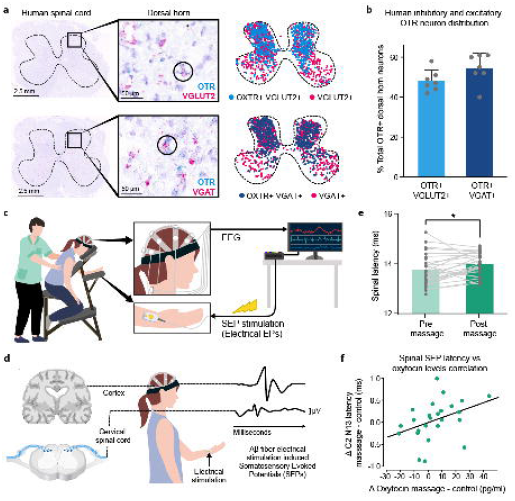
Human spinal OTR circuits mirror mouse mechanisms. a. Chromogenic in situ hybridization (RNAscope) was performed on human lower thoracic to lumbar spinal cord tissue harvested from brain-dead organ-donors (50-80 years old, 2 males 1 female). OTR, VGLUT2 and VGAT probes were used and revealed overlap of OTR expressing cells with both VGLUT2 and VGAT mRNA in the human dorsal horn. b. Quantification of OTR+ neurons among total neurons shows 48.14% of OTR cells express VGLUT2 and 54.43% of OTR cells express VGAT in the human dorsal horn. c. Schematic of participants receiving an electrical skin stimulation before and after a light pressure skin-to-skin back massage, coupled with Somatosensory Evoked Potential (SEP) recording. d. SEP recordings were obtained at the C2 cervical spinal level and at the cortical level near the somatosensory cortex (for supplementary analysis) to record electrical skin stimulation-evoked SEPs and measure their latency before and after massage and endogenous oxytocin release. e. The latency of SEPs at the spinal cervical C2 level was significantly longer (mean = 14.07 ms) after massage compared to before (mean = 13.76 ms). f. A significant positive correlation (Kendall’s τ = 0.343, p = .008, one-tailed) indicates that greater oxytocin increases were associated with larger spinal latency shifts, supporting a modulatory role of oxytocin at the spinal level. Error bars indicate standard deviation.

### Oxytocin–spinal coupling analyses

To examine the relationship between endogenous oxytocin release and spinal somatosensory processing, condition-specific change scores were computed for both measures. For oxytocin, change scores reflected the difference between post- and pre-massage levels, contrasted against the corresponding change in the control session (massage – control). For spinal somatosensory evoked potentials (SEPs), latency change scores were computed at the cervical C2 level by subtracting baseline latency from post-intervention latency at the second post-massage time point (t2), again contrasted against the control session. Associations between oxytocin and spinal latency changes were assessed using Kendall’s tau correlation, selected due to non-normal distributions of hormonal measures. Correlation tests were one-tailed, reflecting the a priori hypothesis that greater endogenous oxytocin release would be associated with increased spinal SEP latency.

Oxytocin–spinal association. A significant positive association was observed (Kendall’s τ = 0.343, p = 0.008, one-tailed), indicating that individuals with greater oxytocin increases during massage also exhibited larger increases in spinal SEP latency after intervention.

### Electrocardiogram (ECG) Measurements

ECG data were collected at two time points: baseline and post-intervention on each session. Post-intervention recordings were taken either after massage (experimental visit) or after completing a questionnaire (control visit). At each time point, ECG was recorded for 2 minutes using the B-Alert EEG/ECG system from BIOPAC (MP160) Acqknowledge. The raw ECG data were processed using EEGLAB in MATLAB. The R-peaks, corresponding to each heartbeat, were detected and inserted into the continuous data using the HEPLAB toolbox. Visual inspection and manual correction of R-peaks was performed. The time between successive R-peaks (interbeat interval, ITI) was extracted and used for subsequent analysis. Heart rate (HR) and heart rate variability (HRV) were calculated using Kubios software (Tarvainen et al., 2014). HRV was quantified based on the ratio of low-frequency to high-frequency components (LF/HF) using parametric autoregressive (AR) modeling, which provides results in normalized units (nu) suitable for short periods of recording (Tarvainen et al., 2014). Heart rate (HR) was baseline-corrected by subtracting baseline values from post-intervention values for each visit. Baseline-corrected HR changes were compared between the massage and control visits using paired-samples t-tests (two-sided, α = 0.05). Analyses were conducted on participants with complete paired data for both visits.

Heart rate. Baseline-corrected HR differed between intervention conditions (paired t-test, t(25) = 2.918, p = 0.007), with a greater decrease during the massage session (mean = −5.23 bpm, SD = 4.56) compared to the control visit (mean = −1.55 bpm, SD = 3.66).

### Human RNAscope studies — Chromogenic in situ hybridization on human spinal cord tissue

Human spinal cord were obtained from four brain-dead organ-donor patients (50-80-year-old) under the approval of the French institution for organ transplantation (Agence de la Biomédecine). The 3 patients (2 men, 1 woman) died from stroke and did not suffered from chronic pain. Body temperature was lowered with ice, and following circulation arrest blood was replaced with a cold perfusion of organ preservation solution. After organ removal for transplantation purpose, a spinal segment from thoracic T8 to lumbar L2 levels was removed in one piece and spinal cord was immediately dissected in ice cold oxygenated HBSS solution. The delay between clamping blood circulation and the spinal cord dissection was ∼3 hours. Tissues were subsequently flash frozen in liquid nitrogen. This short time interval allowed good preservation of the tissue as previously shown58. For subsequent in situ hybridization experiments, frozen spinal cord 12 µm sections were prepared with a cryostat and mounted on SuperFrostPlus (ThermoFisher). Due to high level of lipofuscin fluorescent background in human samples, we performed duplex chromogenic RNAscope in situ hybridization assay (RNAscope™ 2.5 HD Duplex Reagent Kit #322430) as instructed by ACD. The probes used were Hs-OTXR (#422651-C1) in combination with Hs-SCL17A6-C2 (# 415671-C2) or Hs-SLC32A1-C2 (# 415681-C2). Sample quality controls were performed with human specific positive and negative control probes (Hs-UBC1 #310041, Hs-POLR2A #310451-C2, Negative control Dap B #320741). Slides were counterstained with hematoxylin. Images were obtained with an Hamamatsu NanoZoomer Slide scanner system and analysed with QuPath 0.3.2 and Fiji softwares.

## Acknowledgments

We gratefully acknowledge the gift of human tissue from the four donors included in this work and their families. Chromogenic RNAscope on human tissue images collected for this manuscript was performed in the Montpellier MRI imaging Platform. We thank Pierre-François Mery for his help in the human spinal cord dissection. CP was supported by ANR-18-CE16-1190 0002-01 and ANR-19-CE14-0033-01. EB was supported by the Labex ICST ANR grant and the FRC neurodon. HO is a Wallenberg Clinical Scholar. JK was supported by an NIA T32 fellowship “Neuroscience of Aging, Neurodegeneration and Alzheimer’s Disease” (AG055378). MAG was supported by an NIH / National Institute of Neurological Disorders and Stroke (NINDS) K99NS133476 award and the New Jersey Commission on Spinal Cord Research. MB was supported by a Simons Foundation Autism Research Initiative Fellows-to-Faculty award. VEA was supported by NIH-NINDS R01NS119268, NIH-NINDS R01NS124799, NIH-NCCIH/NINDS U24AT011970, the Pew Charitable Trust, the Whitehall Foundation, the Craig H. Neilsen Foundation, the Rita Allen Foundation, the Alfred P. Sloan Foundation, the New Jersey Commission on Spinal Cord Research. YUI was supported by JSPS KAKENHI 20K06467, 23K05612.

---

**Figure S1:**
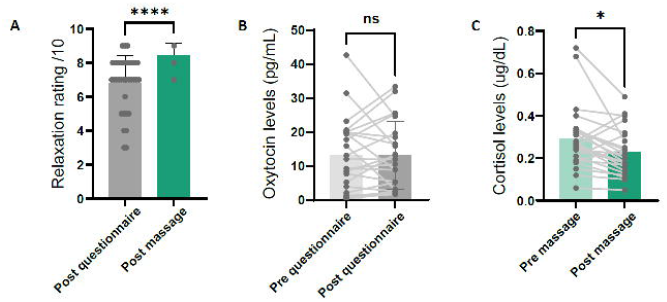
Gentle touch correlates with reduced cortisol and heart rate in humans and induces place preference in mice with neuropathic pain. a. Relaxation ratings (0-10) were significantly increased post-massage compared to pre-massage, confirming the effectiveness of the massage intervention on subjective relaxation state; these data will also be reported elsewhere. b. Salivary oxytocin levels showed no significant change between pre- and post-questionnaire timepoints during the control session, confirming that oxytocin release was specific to the massage intervention. c. Salivary cortisol levels were significantly reduced following massage compared to baseline (Tukey-corrected post hoc comparisons: baseline vs. post: p = 0.047), indicating a physiological relaxation response to massage; these data will also be reported elsewhere. d. Heart rate was significantly more reduced following massage (mean = −5.23 bpm) compared to the control session (mean = −1.55 bpm; paired t-test, t(25) = 2.918, p = 0.007), confirming autonomic relaxation effects of the massage intervention. e. Pleasantness ratings (0-10) collected after the massage session confirmed that all participants rated the back massage as pleasant, validating the positive valence of the tactile stimulation used in this study. f. Mice with established neuropathic pain (Spared Nerve Injury) showed significant gentle brush-induced conditioned place preference, confirming that social touch-like stimulation remains rewarding even in chronic pain conditions.

**Figure S2:**
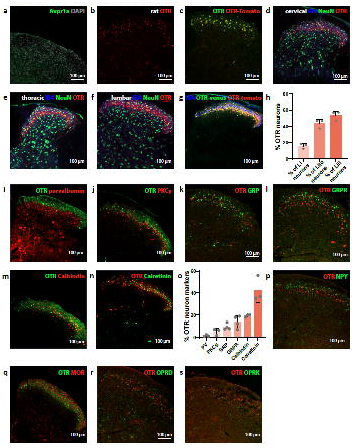
A newly generated OTR reporter OTR-Tomato mouse line reveals the variety of dorsal horn OTR neuron molecular profiles. a. RNAscope using a mouse AVPR1a probe (green) on wild-type mouse spinal cord tissue reveals very low AVPR1a expression levels, supporting the selectivity of oxytocin signaling through OTRs rather than vasopressin receptors in the spinal cord. b. RNAscope using a rat OTR probe (red) on rat spinal cord tissue confirms OTR expression in the superficial dorsal horn, consistent with mouse expression patterns. c. RNAscope using a mouse OTR probe (green) combined with tomato antibody (red) on OTR-Tomato mouse spinal cord tissue confirms that tomato reporter expression faithfully recapitulates endogenous OTR mRNA expression in the superficial dorsal horn. d–f. Immunohistochemistry using tomato (red), NeuN (green), and IB4 (blue) antibodies on OTR-Tomato mouse spinal cord at cervical (d), thoracic (e), and lumbar (f) levels confirms that OTR expression in the superficial dorsal horn is consistent across spinal segments. g. Immunohistochemistry using tomato (red), GFP (green) antibodies, and IB4 (blue) staining on OTR-Tomato; OTR-Venus double reporter mouse spinal cord confirms overlap between the two OTR reporter lines, validating both lines for characterization of spinal OTR populations. h. Quantification of OTR neurons across laminae I, IIo, and IIi as a proportion of total superficial OTR neurons shows that 17% of superficial spinal OTR neurons are located in lamina I, 43% of lamina IIo, and 53% of lamina IIi neurons. i–n. Immunohistochemistry on OTR-Venus spinal cord tissue using antibodies against parvalbumin (i), PKCγ (j), calbindin (m), and calretinin (n), and RNAscope on OTR-Tomato spinal cord tissue using probes against GRP (k) and GRPR (l), revealing molecular diversity of superficial spinal OTR neurons and overlap with established dorsal horn interneuron populations. o. Quantification of the proportions of parvalbumin+, PKCγ+, GRP+, GRPR+, calbindin+, and calretinin+ neurons among total superficial dorsal horn OTR+ neurons, confirming broad molecular heterogeneity of the spinal OTR population. p–s. Immunohistochemistry on MOR-Cherry; OTR-Venus spinal cord tissue (p) and RNAscope on OTR-Tomato spinal cord tissue using probes against OPRD (q), OPRK (r), and NPY (s) reveal minimal overlap (2% or less) between spinal OTR neurons and opioid receptor- or neuropeptide Y-expressing populations.

**Figure S3:**
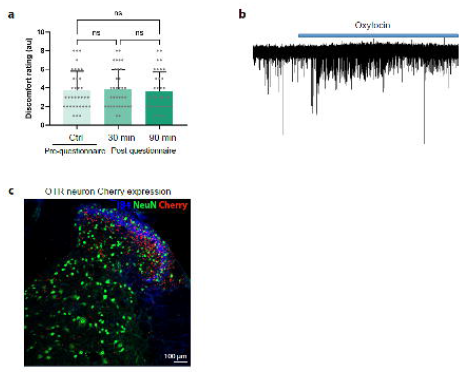
Massage-induced oxytocin release correlates with reduced discomfort ratings of electrical stimulation in humans; oxytocin increases superficial spinal OTR neuron excitability in mice. a. Subjective discomfort ratings of electrical skin stimulation showed no significant change across timepoints during the control questionnaire session, confirming that the reduction in discomfort observed in the massage session was specific to massage-induced oxytocin release and not attributable to habituation or time effects alone. b. Patch-clamp recordings from OTR-Tomato+ spinal neurons show that bath application of oxytocin increases firing probability and action potential frequency in superficial spinal OTR neurons (7/12 OTR+ neurons depolarized), consistent with OT-mediated excitation of this population. c. Immunohistochemistry using tomato (red) and GFP (green) antibodies on OTR-Cre; Ai3; OTR-Tomato spinal cord tissue reveals near-complete overlap between OTR-Cre-driven GFP expression and OTR-Tomato reporter expression in the superficial dorsal horn, confirming that the OTR-Cre line targets the same superficial OTR population characterized in Figure 2.

**Figure S4:**
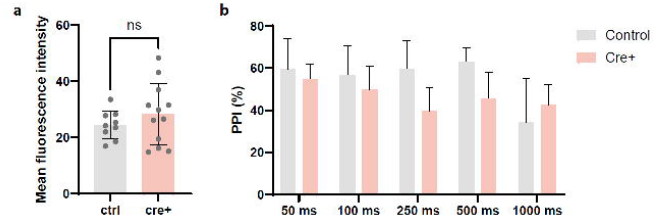
Conditional spinal OTR knockdown is restricted to the injection site and does not affect baseline tactile sensitivity of back hairy skin. a. RNAscope using a mouse OTR probe on cervical spinal cord tissue from control (OTR-flox/flox) and conditional knockdown (OTR-flox/flox::AAV-Cre) mice reveals comparable OTR expression levels between groups at the cervical level, confirming that viral-mediated Cre expression and OTR knockdown are restricted to the lower thoracic injection site and do not affect supraspinal oxytocin receptor signaling. b. Tactile and acoustic prepulse inhibition (PPI) measurements in control and spinal OTR cKO mice show no significant difference between groups, confirming that back hairy skin mechanosensory detection and spinal sensorimotor gating are preserved following lower thoracic OTR knockdown. Together, these controls establish that the loss of gentle brush-induced place preference observed in OTR cKO mice reflects a specific impairment of affective-motivational touch processing rather than a deficit in basic sensory detection.

**Figure S5:**
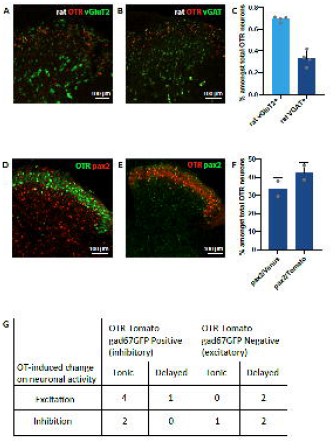
Spinal OTR neurons are a mixed inhibitory and excitatory population differentially modulated by oxytocin. a–c. RNAscope using OTR and VGLUT2 probes (a) and OTR and VGAT probes (b) on rat spinal cord tissue, with quantification (c), reveals that spinal OTR neurons in rats consist of 69% excitatory (VGLUT2+) and 33% inhibitory (VGAT+) populations, confirming the mixed excitatory-inhibitory phenotype of spinal OTR neurons across rodent species. d–f. Immunohistochemistry using GFP and pax2 antibodies on OTR-Venus spinal cord tissue (d) and tomato and pax2 antibodies on OTR-Tomato spinal cord tissue (e), with quantification (f), confirms that 38% of superficial spinal OTR neurons have an inhibitory neurotransmitter phenotype, consistent with RNAscope-based estimates in mice. g. Summary table showing the proportions of tonic and delayed-firing excitatory and inhibitory OTR-Tomato+ neurons whose firing patterns are modulated by bath oxytocin application, revealing a preferential modulatory effect of oxytocin on inhibitory OTR neurons compared to excitatory OTR neurons.

**Figure S6:**
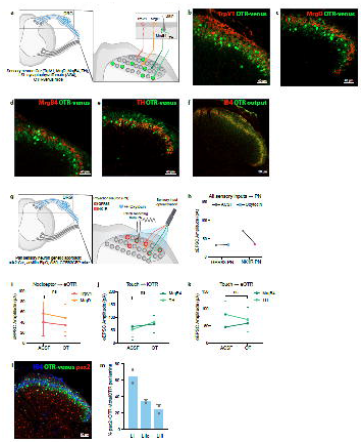
Superficial spinal OTR neurons receive both innocuous and noxious peripheral inputs and project onto spinal output pathways associated with positive and negative valence. a. Schematic of the genetic strategy used to visualize sensory fiber inputs onto OTR neurons: TrpV1-Cre, MrgD-CreER, MrgB4-Cre, or TH-CreER mice were crossed to advillin-FlpO and FPsit reporter mice on an OTR-Tomato background to label specific sensory fiber terminals and OTR neurons simultaneously. b–e. Immunohistochemistry on TrpV1-Cre (b), MrgD-CreER (c), MrgB4-Cre (d), and TH-CreER (e) mice crossed to advillin-FlpO; Ai34/FPsit; OTR-Tomato/Venus reporter lines shows direct apposition of noxious (TrpV1+, MrgD+) and innocuous touch-sensitive (MrgB4+, TH+) sensory fiber terminals onto superficial spinal OTR neurons. f. Immunohistochemistry using GFP and IB4 antibodies on OTR-Cre; Lbx1-FlpO; FPsit spinal cord tissue shows that the axonal outputs of spinal OTR neurons are concentrated in lamina I and lamina IIo, as defined by IB4 staining, positioning OTR neurons to directly modulate superficial output pathways. g. Schematic of the intersectional genetic strategy used to broadly activate somatosensory inputs and record from GPR83+ and putative NK1R+ spinoparabrachial projection neurons with and without bath oxytocin: cdx2-Cre; advillin-FlpO; Ai80; GPR83-GFP mice were used with retrograde labeling from the lateral parabrachial nucleus. h. Quantification of opto-evoked EPSC amplitudes from broad somatosensory input stimulation onto GPR83+ and putative NK1R+ spinoparabrachial projection neurons before and after bath oxytocin application. OT did not significantly affect GPR83+ neuron responses, while it significantly reduced opto-evoked EPSC amplitude in putative NK1R+ spinoparabrachial neurons, suggesting an overall inhibitory effect of OT on avoidance-associated output pathways. i–k. Quantification of opto-evoked EPSC amplitudes in response to noxious inputs (TrpV1+, MrgD+) onto excitatory OTR neurons (eOTRs) (i), and innocuous touch inputs (TH+, MrgB4+) onto inhibitory OTR neurons (iOTRs) (j) and excitatory OTR neurons (eOTRs) (k), before and after bath oxytocin application. No significant differences were observed in these conditions, confirming the selectivity of OT modulation for iOTR responses to noxious inputs and for GPR83+ output pathway responses to touch inputs. l–m. Immunohistochemistry using tomato, IB4, and pax2 antibodies on OTR-Tomato spinal cord tissue (l), with quantification (m), reveals a dorsoventral gradient in the inhibitory-to-excitatory ratio of OTR neurons: inhibitory OTR neurons (pax2+) predominate in lamina I (64.3% of OTR+ neurons) and decrease progressively toward lamina IIi (23.7% of OTR+ neurons), providing an anatomical basis for the lamina-specific modulation of noxious versus innocuous processing by spinal oxytocin.

**Figure S7:**
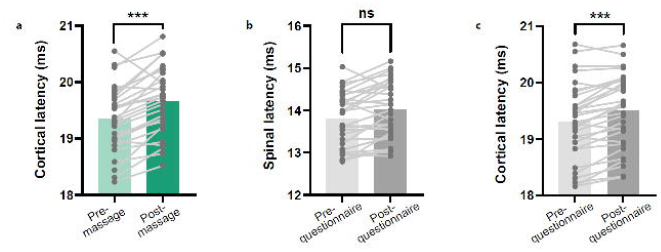
Massage, but not questionnaire control, correlates with increased somatosensory latency; oxytocin selectively predicts spinal rather than cortical latency changes. a. SEP latency at the cortical level was significantly increased following the massage session, consistent with massage-induced modulation of somatosensory processing propagating from spinal to cortical levels. b. SEP latency at the spinal cervical C2 level showed no significant change following the control questionnaire session, confirming that the spinal latency increase observed post-massage was specific to the massage intervention and not attributable to habituation or time-of-day effects. c. SEP latency at the cortical level was also increased following the control questionnaire session, indicating that cortical latency changes are not specific to massage and may reflect broader attentional or temporal factors independent of oxytocin release.

